# Information Spillover in ‘Resting Memory’ and ‘Working Memory’

**DOI:** 10.1101/2025.06.26.661694

**Authors:** Jan Willem Koten, Michael Vogrin, Guilherme Wood, Hans Manner

## Abstract

Our fMRI study investigates the dependencies between brain areas using directed spillover indices estimated from vector autoregressive models that recognize dynamics in the network. Spillover analysis examines how brief neural signal bursts evolve over time and how these dynamic changes can predict past and future brain states across different regions of the brain network. Information spillover was assessed using a test-retest design to estimate the *neural memory capacity* across distinct cognitive states. A dorsolateral prefrontal centered system (DLPFC) demonstrates neural memory capacity at rest, labeled “resting memory”. Resting memory contains roughly 9 times more neurocognitive dependencies (spillover) as the difference in spillover between working and resting brains, suggesting that resting brains exhibit substantial neural memory capacity. The transitioning from “resting memory” to “working memory” is initiated by a right inferior fontal (IFG) centered system which connects to the DLPFC centered system when relevant information is detected in the outside world and also inhibits self-referential feedback in parietal cortices. Spillover between the IFG and DLPFC centered systems facilitate a smooth transition in attention from events that take place outside the brain to (sustained) representations of external events within the brain.

## Main

The term ‘working memory’ yields over 70,000 hits on PubMed, making it impossible to cover all existing models and findings in this introduction. Memory is often categorized into short-term memory, which involves holding information briefly, and working memory, which is the ability to hold information while simultaneously manipulating information [1,2]. The latter is the object of this paper. Classically, working memory was linked to sustained neural firing within the dorsolateral prefrontal cortex [3,4]. Whether frontal regions show “*true sustained activity*” is debatable as electrophysiology has shown that sustained activity is probably an artifact of an averaging procedure [4]. More recently, working memory is viewed as a process in which a short signal burst is transmitted to other regions. This process has been described by Miller and coworkers as: *‘in the constant chatter of the brain, a brief scream is heard better than a constant whisper’*[4]. In this context, it is suggested that short signal bursts (or “screams”) undergo dynamic changes as they propagate through the network[4]. This means that the shape of the signal at the initial time point is not identical to the signal after it has traversed in multiple steps through the network[4,5].

Thus far, several approaches have been employed to unravel how neural information traverse the expanse of neural networks. Among these are structural equation modeling, vector autoregressive models, dynamic causal modeling, and transfer entropy[6,7]. It must be noted that some of these methods are not without their limitations, as elucidated by Ramsey and colleagues [6]. While these methods have improved our understanding of neural networks, they may be less suited to capture the evolutionary changes in signal bursts, over time. In our perspective, “*neural screams*” as they may occur during working memory processes show a remarkable resemblance with “*shocks*” as they occur in economic systems which are described in terms of spillover. The use of econometric models for studying memory in the brain is not ad hoc. Similar to the concepts of long-term and short-term memory in psychology, the terms ‘long memory’ and ‘short memory’ are employed in economics to describe memory processes present in time courses [8,9].

Econometric models have been developed that estimate how shocks are transmitted through networks over time that potentially may also improve our understanding of relationships in the brain. Connectedness measures based on Vector autoregressive (VAR) models assess the distribution of forecast error variation within and between brain regions, after a certain number of periods, originating from neurocognitive shocks initiated in previous time points. This model treats the underlying network as a dynamic system. Recently, Diebold and Yilmaz [10,11] have made substantial advancements to connectedness measures within the realm of financial markets. By determining the directionality of neural connections, the VAR-based connectedness measures can establish whether a brain region is sending or receiving information, a phenomenon known as spillover as introduced in Diebold and Yilmaz [10,11]. By aggregating the spillover effects for each brain region, it is possible to classify a region as a net sender or net receiver of information. An additional advantage of spillover measures is that they can be employed to estimate the total amount of information processed within the network. Contrary to the concept of Granger causality and simple VAR models that are sometimes used in imaging, spillover measures can estimate how much information is processed in feedback loops within a few time periods, which is of advantage as the influential multi component model by Baddeley and Hitch proposes that information is looped between brain areas [12]. Spillover may be interpreted in terms of explained variance in a similar way as R^2^ in the context of regression analysis.

As mentioned, spillover describes how dynamically evolving signals can predict both future and past states. We hypothesize that working memory involves two dynamic copying processes. First, information from the outside world must be duplicated in the brain. Second, this neural representation must be copied to the next point in time, requiring stability; as an unstable representation would vanish quickly. *Neural memory capacity* can be viewed as a process where neural information is repeatedly copied forward. In this framework, the VAR process can be seen as a representation of the “neural copy process”. As neural copies are recursively generated over time, a first-order VAR model may adequately represent the underlying process. However, it may be beneficial to explore connectivity beyond the next time point (lag 1) to include more distant future points. In short, working memory involves a process where attention transitions from a stimulus present outside the brain to a dynamically sustained neural representation of the very same stimulus within the brain.

It is likely that dynamic processes are also present during resting state, as cognitive or emotional processes during rest do not occur randomly, but are most likely influenced by the previous state. Or in the words of Coppola and colleagues: “*We base our approach on the phenomenological observation that any individual experience can be characterized by the experiences anteceding it and the experiences it may lead to*”[13]. In our own words, we hypothesize that as the mind freely wanders from one place to another, it does so in small steps where a neural memory process connects past brain states with present and future states to create the stream of consciousness^[13],[14]^. It is difficult to measure the actual content of the stream of consciousness by scientific means (yet) as it relies on introspection ^[13–15]^. But it is possible to investigate if resting state and working memory exhibit neural memory capacity in a common brain network. Since working memory depends on the conscious processing of information over time, one might expect that the neural correlates of resting and working states exhibit similar neural behavior.

To better understand this phenomenon, we conducted a study in which spillover effects of verbal and spatial working memory data were compared to resting state data. We considered these two tasks because verbal and spatial working memory have been extensively investigated possibly due to the influential multi component model [12]. Participants were asked to hold two spatial positions or two letters in working memory while simultaneously performing a distracting number Stroop task, which required them to identify the physically larger of two simultaneously presented numbers.

Since spillover statistics are inherently positive, we decided to assess images based on reproducibility rather than relying solely on significance testing. For this reason, both resting and working state experiments were measured twice in two independent fMRI sessions. We constructed a connectome from 34 brain coordinates derived from an fMRI meta-analysis of working memory and used it for a spillover analysis [16]. It should be stated that working state coordinates are more reproducible that resting state coordinates which is our main argument to prefer working state coordinates [17–19]. Studies examining the reproducibility of comparisons between two cognitive states known as narrow contrast are scarce. The limited research conducted suggests that the reproducibility of narrow contrasts is low possibly due to the poor contrast to noise ratios [20].

## Results

### Short overview

In the initial step, we will present standard connectivity maps estimated from Pearson correlations encompassing 34 brain regions. Next, we will reduce the connectome to 14 regions and identify the most prominent brain connections associated with neural working memory capacity by subtracting resting-state spillover maps from their working-state counterparts. The resultant hubs in the inferior frontal gyrus (IFG) and dorsolateral prefrontal cortex (DLPFC) are subjected to a detailed analysis. Specifically, we investigate how the working and resting state may show commonalities and differences with regard to neural memory capacity. Next, we will examine whether specific regions can be classified as primarily receiving or primarily sending areas. Finally, we will report the total neural memory capacity of both resting and working brains.

### Results for standard connectivity analysis

The 34 regions of interest that were obtained from working memory-relevant coordinates derived from meta-analysis were subjected to conventional connectivity analysis, resulting in a connectome of 561 paths [16]. A path was considered reproducible at the group level when it was statistically different from zero using a (two-sided) Bonferroni-corrected p-value threshold of 0.05/(561*2) in both the test and retest runs. This procedure is known as conjunction analysis [21]. To further improve the trustworthiness of our results, we only accepted the results of the conjunction analysis when the individual paths showed sufficient test-retest reliability ICC(2,1)>0.4 (**Supplementary Fig. 1**), which is generally considered to be fair [22]. Almost all of the 561 examined paths showed significant results in both working and resting states and are not discussed further. The connectome that resulted when spatial WM was contrasted with resting state is shown in **Fig. 1**. Bilateral hubs in medial systems as well as frontal parietal hubs mainly located in the right hemisphere were reliably detected. A similar contrast for the verbal WM task revealed again a mainly right hemispheric system that was however connected with a language relevant hub on the left superior temporal gyrus (STG) which is roughly equivalent with Wernicke’s area. Furthermore, a path which connected the anterior part of the left intra parietal sulcus (IPS) with the left pre central gyrus (PCG) showed decreased activity compared to resting state in both the spatial and verbal WM tasks. Standard connectivity analysis cannot discern memory relevant from memory irrelevant paths as it does not capture neural memory capacity nor can it provide information on the direction of the information transfer. Hence a spillover analysis was conducted.

**Figure 1:**
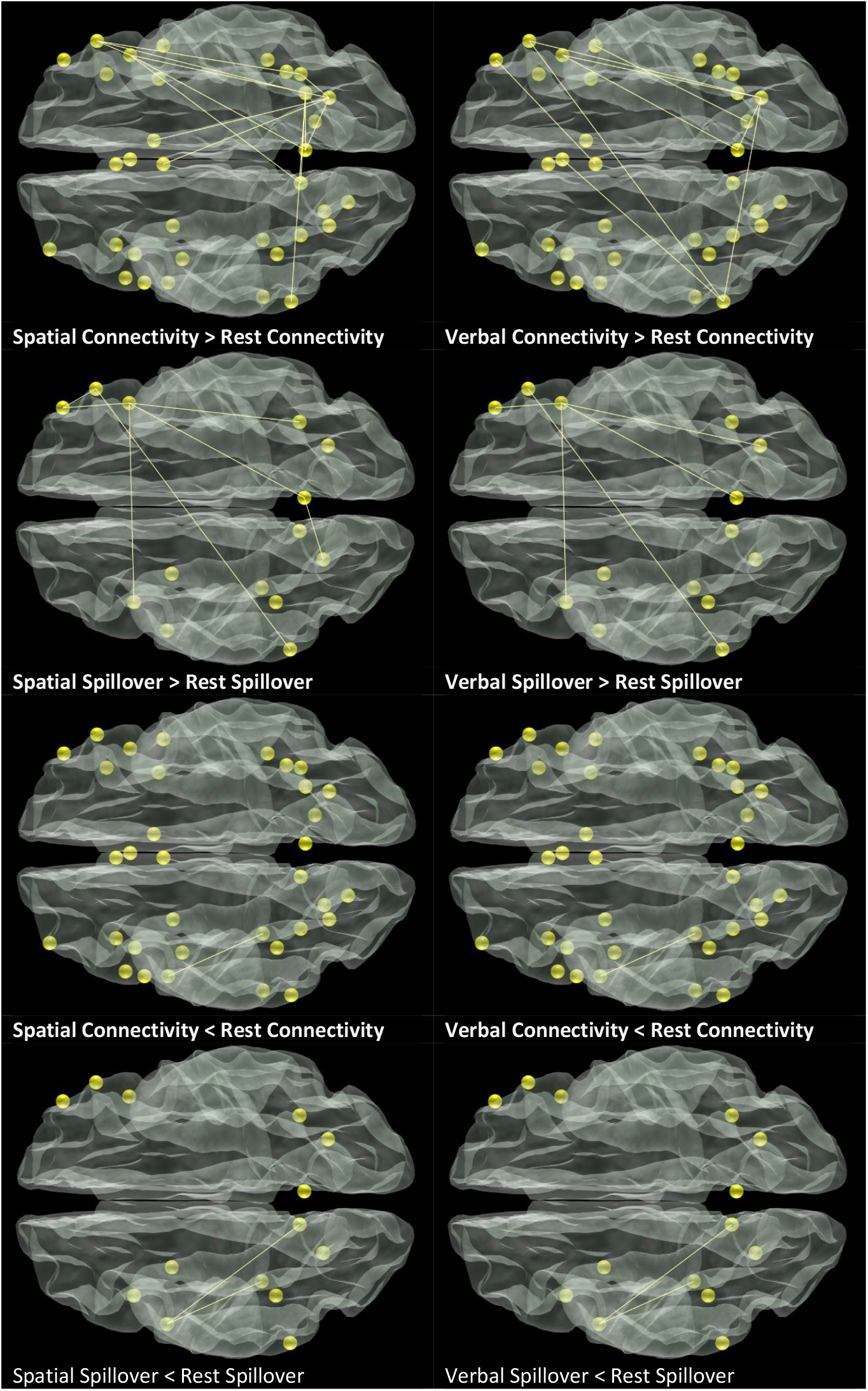
This figure illustrates the connections between brain regions as determined through both conventional connectivity analysis and VAR-based analysis. In order to be included in the results, both the test and retest runs had to meet a Bonferroni-corrected threshold value, and each individual connection had to demonstrate a minimum ICC>0.4.

### Pathwise spillover contrast

The aim of this analysis is to isolate working memory paths that show increased neural memory capacity compared to resting state. The number of ROI’S for the spillover analysis were reduced from 34 to 14 with a filtering procedure described in the method section. Given the stability of the mean spillover maps across different lag orders (**Supplementary Fig. 2-7; Supplementary Table 1**), as well as clear statistical evidence (**methods)**, we focus our analysis solely on the VAR(1) model. The task-related signal averages (**Supplementary Fig. 8-10)** show substantial differences in how brain regions respond to the task, an essential prerequisite for spill-over analysis. The key spillover statistics are reported in **Tables 1 to 3** for working and resting state. The off-diagonal elements show the percentage of forecast error variance explained by neural shocks from other brain regions. The column direction indicates to which area information is sent, while the row direction shows from which area it receives information. Within region estimates reported along the diagonal are substantially higher than between region estimates reported off diagonal irrespective of run or condition. This suggest that a large fraction of neural information stems from the region itself (**Tables 1-3**). Subsequently, the pathwise test-retest reliability for the 196 spillover paths were estimated (**Supplementary Fig. 11**). The average test-retest reliability was 0.68 for spatial WM; 0.63 for verbal WM and 0.53 for resting state data.

**Table 1.**
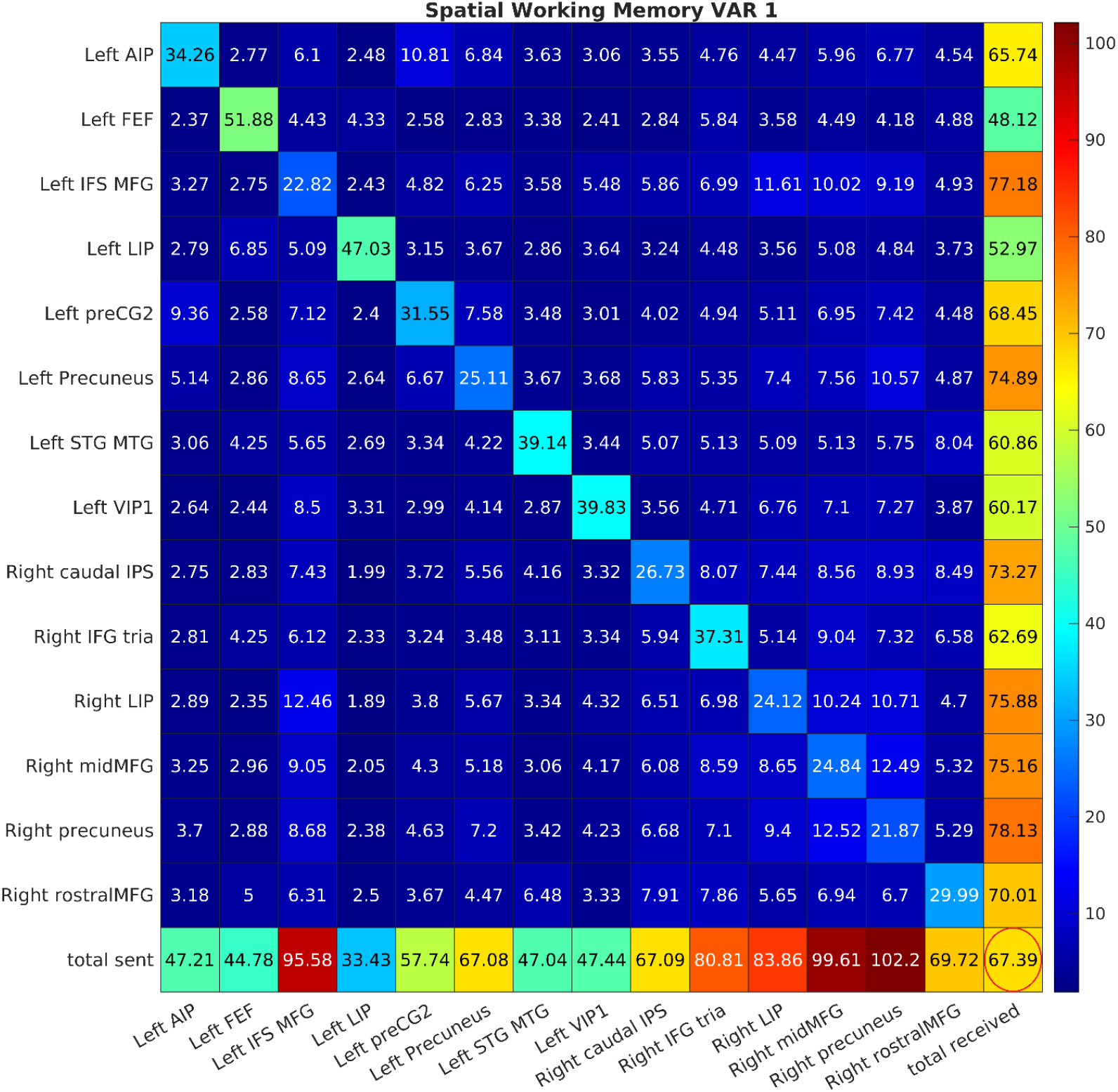
This table reports the grand average spillover statistics obtained from a spatial WM task. The elements on the diagonal represent the amount of forecast error that is attributed to neural shocks as they were generated within the region itself. The off-diagonal elements represent the percentage of forecast error that is explained by neural shocks stemming from other brain regions. The column direction displays to which brain area a specific region is sending the row direction displays from which brain area a region is receiving information. According to established conventions the bottom row represents the sum of the column values excluding the diagonal which represents the total amount of spillover that was sent by a region while the right most column represents the amount of received spillover. The difference between the total spillover sent and the total spillover received is referred as net spillover. The number encircled in red at the bottom right corner represents the total spillover, defined as the sum of the spillover from the off-diagonal elements divided by the total spillover of all elements multiplied by 100.

The mean reliability for complex contrasts was 0.41 for spatial WM-minus-rest; 0.35 for verbal WM-minus-rest and 0.27 for spatial WM-minus-verbal WM. We contrasted the working spillover maps with the resting spillover maps. While the reduced connectome contains significantly fewer connections than the original connectome from which it was derived, we chose to implement a stringent Bonferroni correction based on the larger connectome. This approach helps to avoid circular reasoning, utilizing a Bonferroni p-value threshold of 0.05/(34 ROI * 34 ROI * 2 sides). Furthermore, we required that conjuncted maps exhibited a minimal path wise test-retest reliability of ICC(2,1)>0.4.

The resulting maps shown in **Fig. 1** reveals that the spillover maps show partial overlap with standard connectivity maps. We created a graph illustrating the strength and direction of spillover in **Fig. 2**. Upon initial examination of the maps, it is evident that the right inferior frontal gyrus (right IFG) is a central structure in verbal and spatial working memory tasks as it shows increased spillover with a larger number of frontal parietal regions. The task related signal averages depicted in **Fig. 3** reveal that the right IFG is the first to show activity when subjects are confronted with the initial encoding stimuli that were presented after a period of rest. This prompts the right IFG to direct its “attention” to external stimuli, while other brain regions take longer to respond. Importantly **Fig. 2** suggests that the right IFG reproducibly initiates action in the inferior frontal sulcus/medial frontal gyrus (IFS/MFG), a region that is considered equivalent to the dorsolateral prefrontal cortex and is traditionally seen as a critical area for working memory. In the case of spatial working memory, a feedback loop between the right IFG and the left IFS/MFG is established. Furthermore, increased spillover between the right IFG and the right rostral aspect of the MFG as well as the right pre cuneus was observed for both the verbal and spatial WM. While there was spillover between the right IFG and right lateral aspect of the IPS in the case of spatial memory this was not the case for verbal memory. Furthermore, the IFG may play a role in suppressing self-referential parietal feedback loops. In the case of spatial WM self-referential feedback loops were suppressed in the right hemisphere and included the lateral aspect of the IPS the precuneus as well as the caudal aspect of the IPS. The right middle part of the MFG that showed signs of self-suppression itself also caused self-suppression in the left superior temporal gyrus/medial temporal gyrus. During the verbal working memory the IFG suppressed self-referential brain activity in the ipsi lateral caudal aspect of the IPS. The narrow contrast spatial WM minus verbal WM was considered less relevant and is discussed in **Supplementary Fig. 12**.

**Figure 2:**
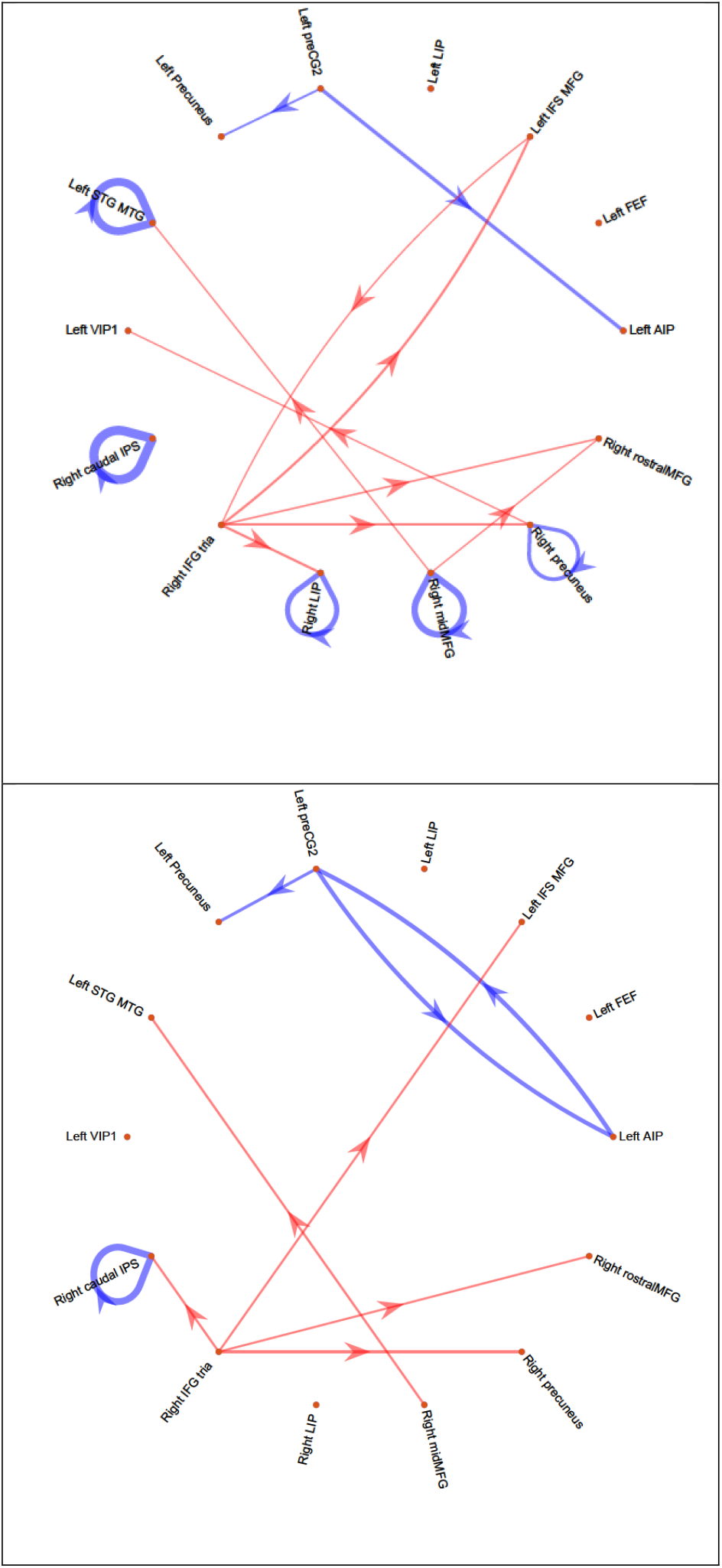
These graphs provide a comprehensive overview of the spillover contrast working versus resting state. Top, the contrast Spatial WM versus Resting state, Bottom the contrast Verbal WM versus Resting state. Red arrows indicate a relative increase in spillover compared to resting state, while blue arrows signify a relative decrease. The line thickness corresponds to the magnitude of the spillover difference between working and resting states. To be included in the results, both the initial test and subsequent retest runs had to exceed a Bonferroni-corrected NHST threshold value, and each individual connection had to display a minimum test-retest reliability of ICC(2,1) > 0.4 for the contrast working-minus-resting state.

**Figure 3:**
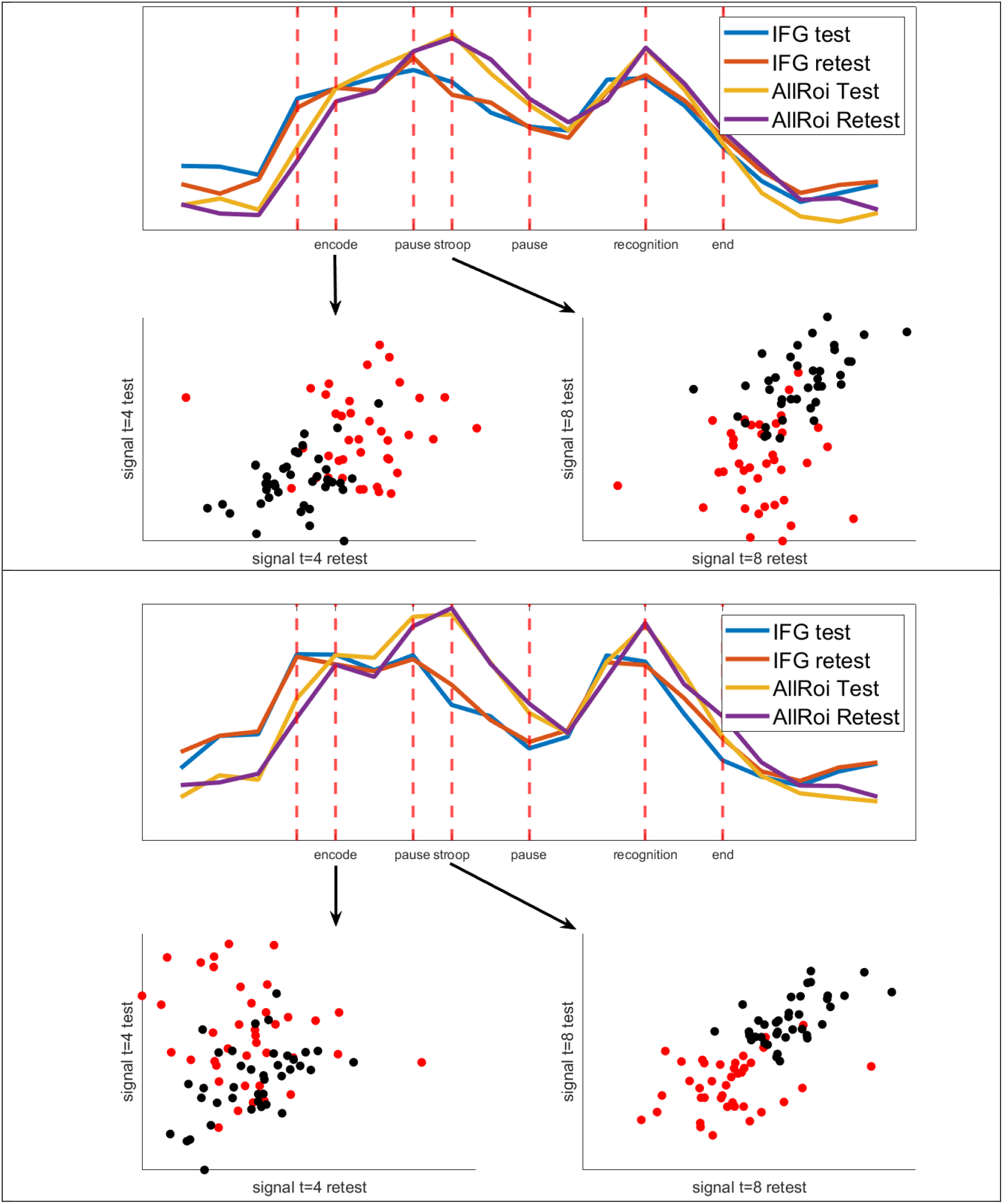
This figure displays the task-related signal averages for the right IFG and the grand mean signal average of 13 remaining brain regions, excluding the IFG. The mean signal was extracted per subject at the timepoints corresponding to the encoding phase (t=4) and the Stroop task (t=8). IFG signals are represented by red dots, while signals from the rest of the brain are shown in black. Top: This image presents the effects observed in the visual working memory task. Bottom: This display highlights the effects from the verbal working memory task.

### Neural memory capacity of IFG and DLPFC centered systems

In our previous analysis we isolated the IFG and DLPFC as potential regions of interest by subtracting resting state maps from working state maps. While these narrow contrasts are helpful to determine regions of functional interest, they also reflect a brain state that does not exist in the real world as a specific brain area cannot be in a working and resting state condition simultaneously. To gain a deeper understanding of the connections established by the IFG and DLPFC with other regions, we focused solely on visualizing the relationships among regions directly connected to the IFG or DLPFC, omitting those that were not linked to these regions. It is not meaningful to test spillover against zero since spillover is always positive. Hence, we implemented an alternative albeit very strict threshold criterion. Specifically, we mandated that regions display a minimum average spillover of 5 and also demonstrate a robust test-retest reliability with an ICC>0.6, indicating a good level of consistency in repeated measurements [22]. Given the absence of established thresholds for assessing spillover effects in the literature, we draw on the guidelines formulated by Funder and Ozer [23]. They indicate that an effect size of r = 0.2 equals a medium effect in correlation studies. In this context, a spillover of 5 can be interpreted as explaining 5% of the variance, which approximately aligns with a medium-sized effect, as 0.2^2^ explains 4% of the variance. Applying these threshold criteria, we discovered that the IFG system shows mainly reliable spillover during working state conditions, while the DLPFC system exhibits reliable spillover during resting and working state conditions. Furthermore, we discovered that both the IFG and the DLPFC system exhibit spillover with a common set of frontal parietal brain regions during working state.

An inspection of the IFG and DLPFC systems during resting state (**Fig. 4**) reveals that the right IFG only shows spillover with spatially neighboring regions including the middle and rostral aspects of the MFG. In contrast, the DLPFC is involved in bilateral fronto-frontal feedback loops and bilateral fronto-parietal feedback loops. The fronto parietal spillover system included the cuneus bilaterally, the lateral aspect of the IPS bilaterally as well as the left anterior aspect of the IPS. In addition, fonto-frontal spillover included the ipsilateral PCG and the contra lateral middle aspect of the MFG. **Figure 4** suggests that DLPFC is a central station that links information streams among brain homologues. It is important to notice that the reported relations found for the DLPFC system were mostly feedback loops with neural memory capacity suggesting that these regions circulate neural information that emerges from within the resting brain itself.

**Figure 4:**
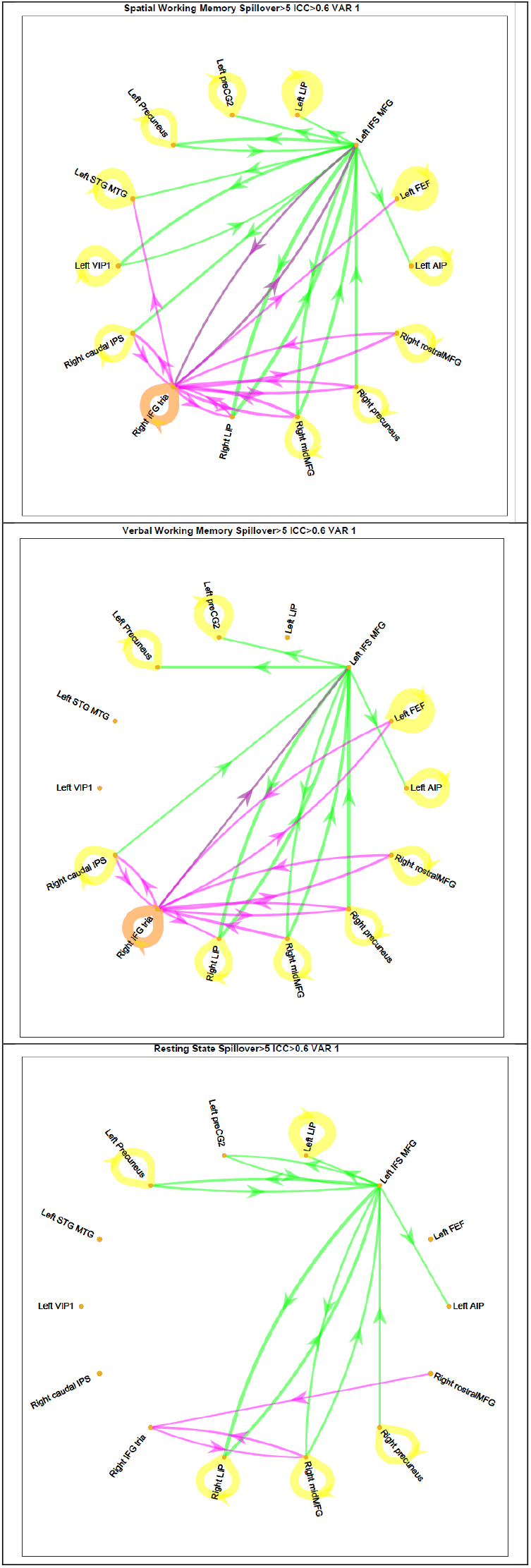
These graphs illustrate the connectivity between brain regions during spatial WM (top), verbal WM (middle), and resting state (bottom) conditions. Pink arrows represent an IFG-centered network, green arrows indicate a DLPFC-centered network, while yellow arrows indicate within region feedback. The line thickness corresponds to the magnitude of the spillover. In order to be included in the results, the average spillover from test and retest runs needed to be above 5, with each individual connection having a minimum test-retest reliability ICC(2,1) > 0.6.

An examination of the IFG and DLPFC systems during working state shows that the IFG experiences increased spillover compared to rest, while the DLPFC displays a network of spillover that closely resembles the pattern seen at rest. The reported relations between the DLPFC and the bilateral cuneus and bilateral LIP as observed during resting state were also present during verbal and spatial working memory suggesting that working memory ‘uses’ neural connections that are also present during resting state (**Fig. 4**). It should be mentioned that not all feedback loops were fully established when the brain was engaged in language, while additional loops with left frontal areas were evident when the brain was engaged in spatial tasks (**Fig. 4**). In short, fronto parietal DLPFC connections are most likely employed to process information that either emerges from the brain itself or information that emerges from the outside world.

During working state, we saw a considerable expansion of the IFG system compared to rest including feedback loops in the same hemisphere involving brain regions such as the MFG, precuneus, and aspects of the IPS (**Fig. 4**). In the opposite hemisphere, the most significant feedback loop was identified in the IFS/MFG (DLPFC). The expansion of the network that is observed when working memory conditions are compared to resting state conditions is a response to the specific demands of working memory tasks. Given the IFG’s role as an early responder to external stimuli, it likely plays a role in allocating frontal parietal resources that may be engaged in attention towards stimuli outside of the brain (**Fig. 2**) that partly overlap with the DLPFC system. We tried to shed light on the partly overlapping nature of the IFG and DLPFC system through an investigation of the task related signal averages and discovered that several phases could be discerned (**Supplementary Fig. 9-10**). While the IFG is the first area to respond to the incoming stimuli, it is important to note that the DLPFC also plays a role as an early responder. The DLPFC may take over the executive function of the IFG when the signal strength of the IFG decreases during the encoding phase (**Supplementary Fig. 9-10**). This transition ensures that executive functions that respond to external stimuli are integrated with executive systems that manage sustained brain activity mainly found in the left precuneus. Importantly the precuneus is an area that responds late to the incoming stimuli (**Supplementary Fig. 9-10**) and is managed by the IFS/MFG (**Fig. 4**). In summary, the IFG system plays a role in transferring stimuli from the outside world into the brain. The DLPFC system manages neural representations of these external stimuli within the brain and activates superior parietal systems that may participate in holding information over time.

### Sending and receiving brain areas

By summing the pathwise spillover estimates in **Table 1-3** column-wise, one can calculate the total information sent; summing row-wise gives the total information received. The convention is to exclude the diagonal entries from these sums. By subtracting the received spillover from the send spillover, it is possible to characterize a region as a net sender or a net receiver. We estimated the test-retest reliability of the 14 net effects under study (**Supplementary Table 2**). The average test-retest reliability obtained from an ICC(2,1) model was 0.66 for the spatial WM; 0.62 for verbal WM and 0.46 for resting state data. The mean reliability for complex contrasts was 0.51 for spatial WM-minus-rest; 0.44 for verbal WM-minus-rest, and 0.31 for spatial WM-minus-verbal WM. As net spillover values range between positive and negative values, we subjected net spillover to significance tests and required that both the test and retest runs should reach a Bonferroni corrected p-value. In addition, contrasts should be detected with sufficient test-retest reliability ICC(2,1)>0.4 and are reported in **Supplementary Table 3**. When contrasts were tested against zero, the rest, verbal and spatial conditions exhibited positive net spillover in the right precuneus. Additionally, the spatial and verbal tasks showed positive net spillover in the right medial frontal gyrus while they also shared negative net spillover in the left anterior and ventral part of the IPS. In addition, we tested spatial WM and verbal WM against resting state. In this case the right IFG pars triangularis (Broca’s homologue) changed from a taking into a giving brain area when the condition changed from resting to working state.

**Table 2.**
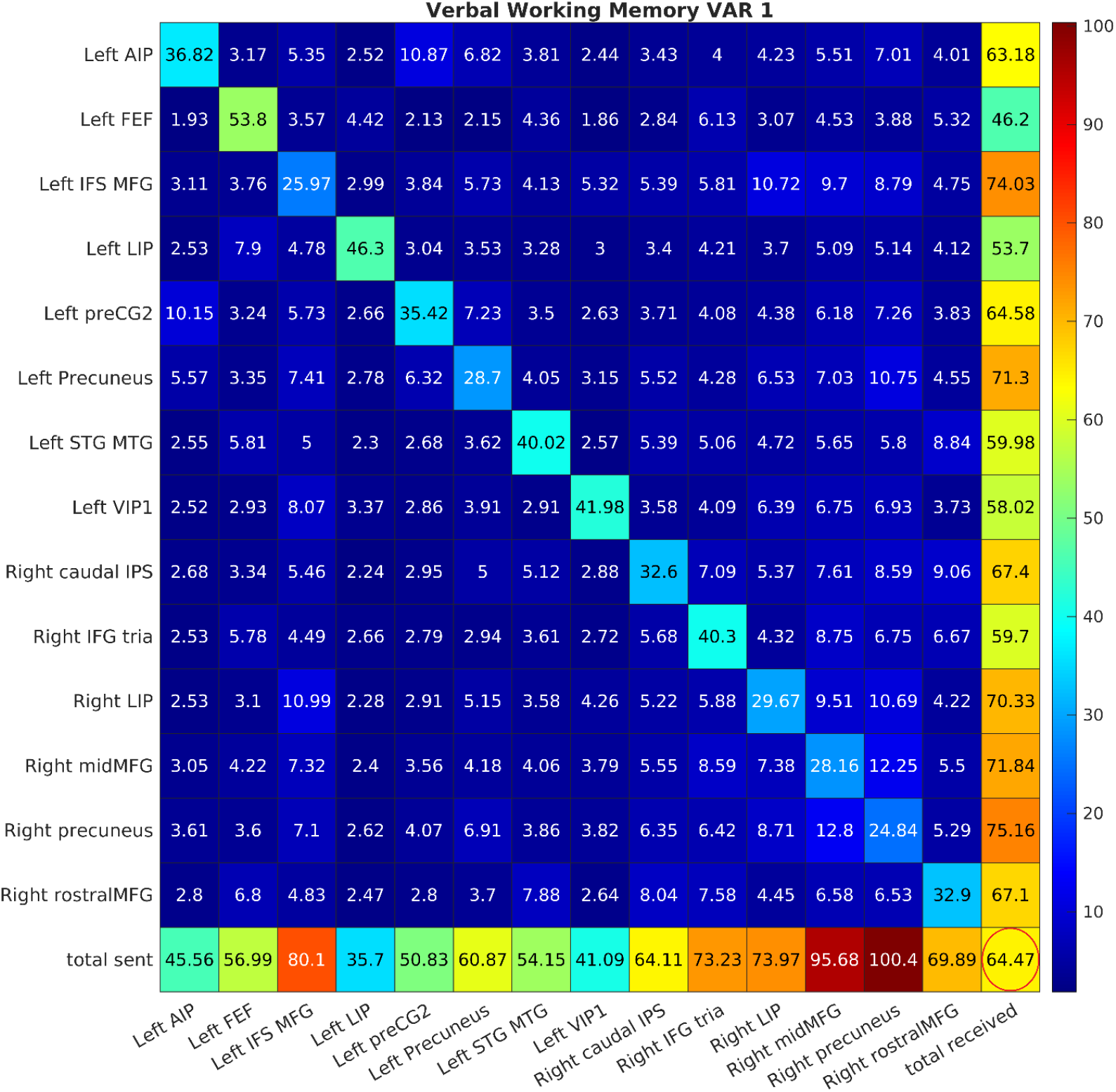
This table reports the grand average spillover statistics obtained from a verbal WM task. The elements on the diagonal represent the amount of forecast error that is attributed to neural shocks as they were generated within the region itself. The off-diagonal elements represent the percentage of forecast error that is explained by neural shocks stemming from other brain regions. The column direction displays to which brain area a specific region is sending the row direction displays from which brain area a region is receiving information. According to established conventions the bottom row represents the sum of the column values excluding the diagonal which represents the total amount of spillover that was sent by a region while the right most column represents the amount of received spillover. The difference between the total spillover sent and the total spillover received is referred as net spillover. The number encircled in red at the bottom right corner represents the total spillover, defined as the sum of the spillover from the off-diagonal elements divided by the total spillover of all elements multiplied by 100.

**Table 3.**
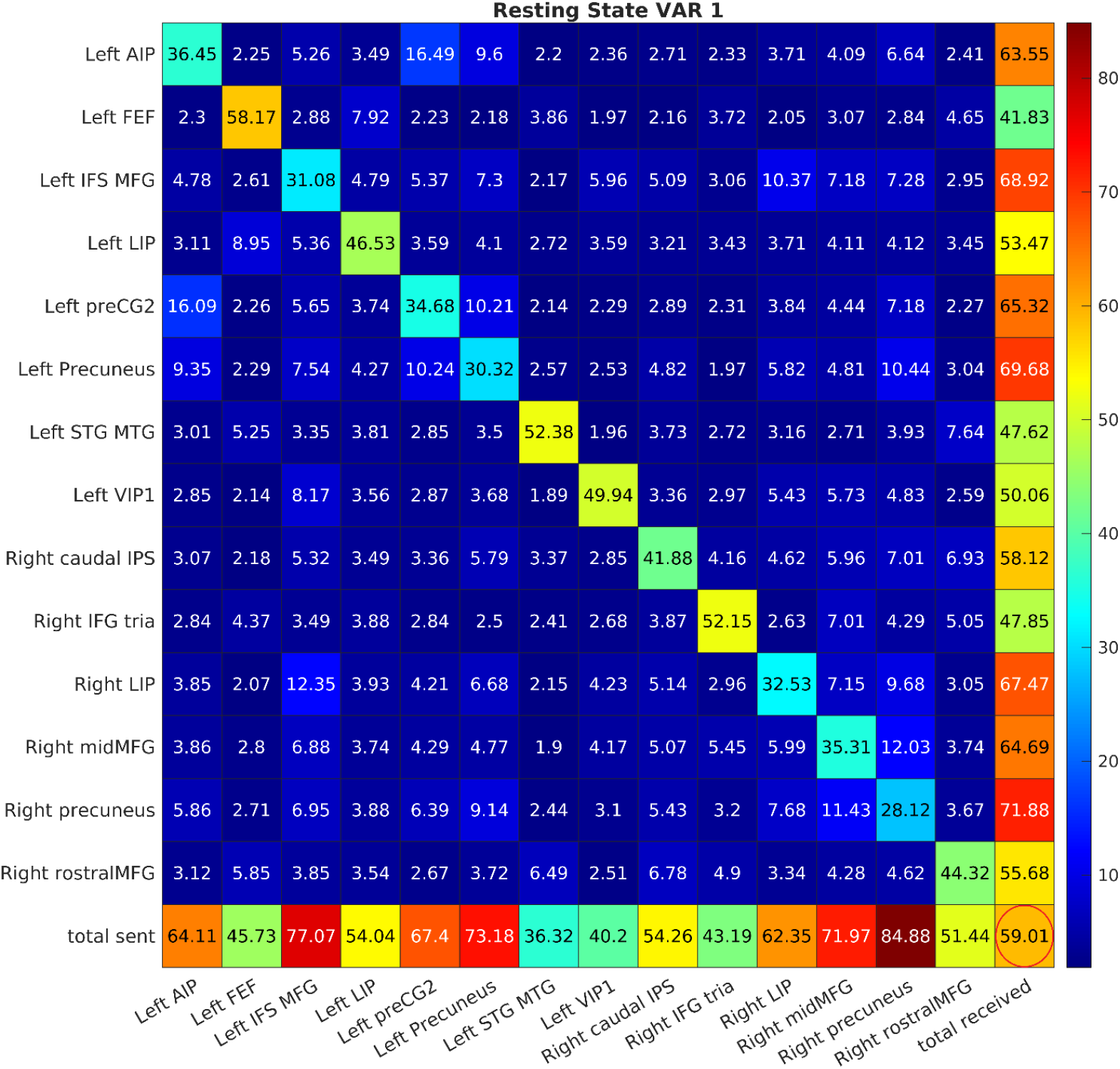
This table reports the grand average spillover statistics obtained from a resting state condition. The elements on the diagonal represent the amount of forecast error that is attributed to neural shocks as they were generated within the regions themselves. The off-diagonal elements represent the percentage of forecast error that is explained by neural shocks stemming from other brain regions. The column direction displays to which brain area a specific region is sending the row direction displays from which brain area a region is receiving information. According to established conventions the bottom row represents the sum of the column values excluding the diagonal which represents the total amount of spillover that was sent by a region while the right most column represents the amount of received spillover. The difference between the total spillover sent and the total spillover received is referred as net spillover. The number encircled in red at the bottom right corner represents the total spillover, defined as the sum of the spillover from the off-diagonal elements divided by the total spillover of all elements multiplied by 100.

### Resting load

The relative amount of neural communication between brain areas was defined as the global amount of received and send information relative to the total amount of information which is equivalent with total spillover. We observed a value of 59 for resting state with ICC(2,1)=0.57; 65 for verbal WM with ICC(2,1)=0.75; 67 for spatial WM with ICC(2,1)=0.71. A conjunction analysis that tested the verbal condition against the resting state condition resulted in p value of 1.29e^-4^ that was traced with an ICC(2,1) = 0.38. The same procedure for the contrast spatial WM versus resting state resulted in a p-value of 1.70e^-8^ with an ICC(2,1) = 0.29. While the conjunction analysis of narrow contrasts shows high reproducibility at the group level the ICC estimates suggest that one should restrain from drawing conclusions at the level of the single subject. The between region communication increased by 9.25% when verbal working memory was compared to resting state while this number was 14.21% when spatial WM was compared the resting state. We estimated the neural load of the resting state condition labeled “resting load” by comparing it with the difference between resting and working state which reflect the “working load”. The resting state condition as such contains 10.81 times more neural load among regions than the difference between verbal WM and resting state. In analogy resting state contains 7.04 more neural load than the difference between spatial WM and resting state.

### Map similarity

We demonstrate that resting state and working state connectomes, obtained through conventional Pearson correlation analysis and spillover analysis, are highly correlated. This relationship is consistently observed not only when assessing standard connectivity and spillover connectomes independently, but also, though to a lesser degree, when comparing pearson connectivity maps to spillover maps. Notably, the correlation between spillover maps and conventional correlation maps is weaker than the correlation observed within their respective categories. For additional details, please see **Supplementary Text 1** with the accompanying **Supplementary Fig. 13-18** and **Supplementary Table 4**.

## Discussion

We hypothesize that working memory processes constitutes of two essential copying steps. First stimuli present in the outside world need to be copied into the brain, second the neural representations of the outside world need to be preserved over time. We tested these simple ideas by means of VAR-based spillover measures that allow to assess how dynamically evolving information is processed in the brain over several time steps. We discovered that copying external events into the brain is partly an IFG guided process while copying neural representations of those external events to the next point in time is partly a DLPFC guided process. We showed that the behavior of the DLPFC centered system is similar during resting and working state. As the spillover captures neural memory capacity, we propose that the DPLFC is possibly engaged in memory processes during rest that may constitute to the (conscious) processing of self-generated information [13].

We propose that resting memory may exist for two key reasons. First, our spillover analysis indicates that the resting memory system shares significant anatomical overlap with a dorsolateral prefrontal cortex centered network that was shown to be active during working memory tasks. Second, both the resting memory system and the working memory system exhibit neural memory capacity. One explanation for this phenomenon may be that even during resting state memory plays a role as the brain is inherently processing information from its own past. While the wandering mind wanders through the “mind space”, some kind of narrative is constructed during the wandering process by linking current cognitive events to past and future brain states [13,14]. This process requires some kind of neural memory capacity that - in contrast to working memory - is not focused on sensory information stemming from the outside world but rather focused on information that is generated by the brain itself. Since there is no nomenclature to describe this memory process, we will refer to it as “resting memory” in contrast to the well-established “working memory”.

The spillover memory analysis depicted in **Figure 4** shows that the DLPFC-centered system has the capacity to copy information to the next point in time during resting and working states. This indicates that the DLPFC system holds information that is either generated by the brain itself (resting state) or originating from the world outside of the brain (working state). The DLPFC exerts a significant influence on a broader network (**Figure 4**), encompassing frontal and parietal areas such as the middle aspect of the medial frontal gyrus, several parts of the IPS and cuneus.

It was once believed that brain regions that show “true sustained” brain activity capture WM related processes. It is tempting to interpret regions exhibiting sustained brain activity, such as the left precuneus (**Supplementary Fig. 8-10**), as engaging in a process where this region copies information internally. However, the spillover data in **Tables 1-3** indicates that only ∼ 30% of the information processed in the left Precuneus stems from the region itself. This suggests that the capacity to maintain within region feedback loops, potentially involved in memory processes, might also depend on spillover interactions with other areas such as the DLPFC (**Figure 4**).

Whether regions show “true sustained activity” is debatable as electrophysiology has shown that sustained activity is probably an artifact of an averaging procedure [4]. Our results are more in line with recent views which claim that memory relevant information is circulated in a cascade of frontal parietal loops which main hubs gradually shift as the memory process commences over time. The actual memory signal is altered when it passes through the frontal parietal cascade structure [4,5]. As an example of this process, both right and left hemispheric brain areas receive input from the IFG and DLPFC, indicating that during the early stages of memory encoding, various regions may transform IFG-guided external information into internal representations, potentially guided by the DLPFC. Data in **Figure 2** shows a decrease in IFG involvement during encoding, while DLPFC-guided parietal systems become more active in maintaining information amid distractions (**Supplementary Fig. 8-10**). It’s important to note that the signal gradually changes over time as it traverses the network, which aligns with modern notions of working memory^[4],^[5]. for instance, the DLPFC receives input from early responders like the right LIP, right precuneus, and right middle MFG, and subsequently sends this information to the left precuneus with a significant time delay[3]. The lagged time course profile is not just a duplication of the DLPFC but has its own distinct characteristics (**Supplementary Fig. 8-10**).

The DLPFC centered system can hold information that stems from the outside world when it is receiving input from the IFG centered system. The IFG centered system may fulfill three functions. First, it detects changes in the world outside of the brain that are memory relevant. Second, it initiates neural processes in the DLPFC that are related to neural memory capacity. Third, it suppresses within region feedback loops occurring during resting states.

Evidence that the IFG copies information that originates from the world outside of the brain to the world inside of the brain stems from the task related signal averages (**Figure 3 and Supplementary Fig. 8-10**). These averages show that the IFG is the first brain area that responds to incoming external information after a period of cognitive rest. This is further corroborated by analyses focusing on the early signal expression of the IFG at the single subject level (**Figure 3**).

The narrow contrast between working and resting states provided in **Figure 2** revealed that the IFG is both driving activity in the left DLPFC and suppressing parietal feedback loops associated with self-referential thinking during rest. The transition from rest to work is not just about the engagement towards external stimuli but also about the suppression of internal, self-referential processes. One possible reason for this suppression is to free neural resources such that they can be used to process incoming external information. Recently, it has been suggested that inhibition is one of the main functions of the frontal systems during working memory processes [4]. It is not impossible that the observed inhibition is related to alpha and beta bursts that are associated with inhibitory processes [4]. In short; working memory may possibly emerge when attention systems designed for the outside world are blended with attention systems tailored for the inside world [3].

Our correlation analysis given in **Supplementary Table 3** proofs that spillover maps show a high level of similarity across resting and working state conditions, and also exhibit some resemblance to standard connectivity maps. While we have demonstrated that differences between working and resting conditions exist it is important to realize that the overall connectedness of the brain is rather stable irrespective of the cognitive content that is processed. Transitioning, from one state into another is a rather subtle process. Notwithstanding these observations we would like to point out that the cognitive load increased with roughly 10% when working memory was compared to resting memory. The resting brain contains roughly 9 times more total spillover information as the difference between working and resting state. The high neurocognitive load of the resting brain labeled “resting load” suggest that the brain is indeed in a condition which we labeled “resting memory”. While it may be challenging to assess the content of the stream of consciousness as it may reside in resting memory through scientific means [13,15], we would like to highlight a tradition in French thought where humans can imagine a state in which the brain is not receiving any information from the senses. A state in which the brain can only observe itself and by doing so discovers that there is an conscious agent that is thinking [24,25]. We tentatively propose a loose analogy between the self-reflective mind and the self-referential nature of temporal processes in the brain.

## Limitations

While we took care to ensure that our results are reproducible within our sample through conventional intra-class correlation analysis employed at the group level, this does not automatically imply that the results are reproducible between samples composed of distinct individuals. In addition, it is inherently impossible to estimate the test-retest reliability of resting state time courses due to the unsynchronized nature of resting state brain behavior. Further research is needed to establish that spillover analysis is a meaningful method to analyze fMRI time courses. Finally, it was beyond the reach of this paper to discuss the neural correlates of resting and working memory at the single subject level[13].

## Methods

### Task and participants

Prior to scanning, participants were trained on the tasks until they performed confidently. Individuals had to hold two letters or two spatial positions in working memory while executing a number Stroop task. During the encoding phase, which lasted 2.5 seconds, participants memorized a set of two letters or two spatial positions. In the distraction phase, also lasting 2.5 seconds, they performed a number Stroop task. Finally, during the recognition phase, which also lasted 2.5 seconds, participants pressed the button with their right index finger when a newly presented item matched the previously presented set, and used their left index finger when the items did not correspond. The encoding and distraction phases were separated by a pause of 1.2 seconds, while the distraction phase and the recognition phase were separated by a pause of 3.7 seconds. The number Stroop tasks consisted of two simultaneously presented Arabic numbers in different physical sizes. Participants were required to identify the larger font size and use their right finger when the larger number was on the right side of the screen, and vice versa. The overall length of the task was 10 repetition times which equals 12.4 seconds. The tasks were separated by jittered resting conditions that lasted ∼12.8 seconds. Details of the task related signal average are given in figure 3. We required that individuals did not make more than 6 wrong responses out of 96 responses (93.75% correct) for both the spatial and the verbal working memory. In total 22 females and 17 males out of 67 individuals survived the selection procedure. The mean age of the population was 41.54 years with a standard deviation of 16.65 which is roughly representative for the Austrian population (https://www.statistik.at/). We estimated the mean response time and the test-retest reliability for the experimental conditions. Results are discussed in the **Supplementary text 2**.

### MRI methods and data preprocessing

MRI scans were performed on a 3 T Siemens Magnetom Skyra (Siemens Medical Systems, Erlangen, Germany) equipped with a 32-channel head coil. Structural images were obtained by means of a 3D-MPRAGE sequence (176 slices per slab, FOV = 256 mm, TR = 2530 ms, TE = 2.07 ms, TI = 900 ms, Flip angle = 9°, voxel size = 1 mm isotropic). Functional imaging data were obtained using a Siemens Grappa parallel acquisition scheme with pat factor 2; using following parameters Flip Angle 72 degrees, TR = 1240 ms, TE = 30 ms. Volume dimensions were 64*64*23, with voxel resolution 4*4*4 mm with a gap of 10%. In total 488 volumes were obtained per run per task while 280 volumes per run were obtained for the resting state condition. The test and retest run were separated by a pause of minimally one hour outside the scanner. The grey and white matter were segmented using FreeSurfer [26]. Subsequently, functional data were brought into 2D FS average space using spherical alignment procedures as available in FreeSurfer. The co registrated grey matter time courses were brought into CleanBrain[27] and subsequently detrended, denoised and corrected for head motions within a GLM framework. The noise regressors consisted of 5 “white matter” components and 5 “ventricle” components that were extracted from 3D space using FreeSurfer. The signal trends were obtained from the grey matter time courses through the application of the SPM low pass filter (128seconds) that was embedded into CleanBrain[27] The motion regressors included the first 2 principal components of the head motion data. We did not extract brain regions from brain activation maxima as this may induce circularity. Instead, 34 3D MNI coordinates that are believed to be essential for working memory were taken from a meta-analysis and brought into 2D FS average space [16].

### Creation of Patches of Interest and Extraction of Grey Matter Time Courses

We did not use spatial smoothing as a preprocessing step. A recent study that investigated the use of spatial smoothing as a preprocessing step in ROI based functional connectivity analysis concluded: *“Spatial smoothing has complex effects on the structure and properties of the networks, including possible over-emphasis of strong, short-range links, changes in the identities of hubs of the network, and decreased inter-subject variation. The ROI approach already includes averaging, independent of spatial smoothing. Therefore, there is no specific reason for applying spatial smoothing”* [28]. Working memory experiments were used in this study to investigate the functional connectivity of frontal parietal systems. Instead of creating patches of interest based on activation maxima potentially present in our own sample, which could cause circularity, we selected 34 MNI coordinates of interest from a recently published meta-analysis focusing on the executive aspects of working memory. These coordinates were brought into FS average mesh space using the procedure described on the FreeSurfer homepage. In short, the 3D coordinates of the ALE meta-analysis were brought into 3D FS average space using the “fslregister” command. Then, the co-registered coordinates were projected onto the FS average mesh using the FreeSurfer “mrivol2surf” command. The relevant mesh elements were brought into MATLAB® format for further preprocessing using the FreeSurfer/MATLAB® command “mri read.” A circle with a diameter of 8mm was drawn around the relevant mesh element using the MATLAB® “surfstat” command SurfStatROI (https://math.mcgill.ca/keith/surfstat/). The resulting vertices were aggregated into a patch of interest (POI), which was used to extract the grey matter time courses of the individuals under study. All the mesh time courses of a specific POI were averaged and used to estimate the connectivity and spillover among nodes, as well as the test-retest reliability. We used a diameter of 8mm which is roughly twice the size of our voxels. This approach mimics to a certain extent the determination of the ideal FWHM of a smoothing kernel on the 2D surface that is believed to be around 8mm[29].

### Conventional connectivity analysis

The 34 time courses were corrected for serial correlations before conventional connectomes were estimated using conventional Pearson correlations. The average autocorrelation functions presented in **Supplementary Table 5** suggest that an ARMA(1,1) filter that was performed on every single time course removes autocorrelations sufficiently [30].

### General linear model

We did not conduct a standard GLM brain activity analysis due to its potential for producing misleading results within the context of a spillover analysis. VAR models assume that time courses in brain activity maps exhibit differential expression and ideally also that time courses are shifted along the time dimension. The commonly used GLM, on the other hand, makes the assumption that time courses should conform to a predictor function that does not account for individual variations in time course behavior. Details of spillover analysis are given below.

### Mathematical Background of Spillover

An attractive approach for measuring spillovers based on reduced-form vector autoregressive models has been proposed by Diebold and Yilmaz [10,11], who applied this approach to measure information spillovers in financial markets. Let *y*_*t*_ = [*y*_1,*t*_ …, *y*_*n,t*_] ′ be a vector valued time-series with the individual series representing n regions of interest in the brain. The reduced form VAR(p) model is given by

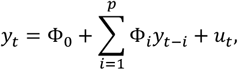

where Φ_0_ is a n-vector of intercepts, Φ_*i*_are *n* × *n* coefficient matrices, both sets of unknown parameters to be estimated by ordinary least squares, and *u*_*t*_is an n-dimensional error representing new information/shocks entering the system. The error has mean zero and covariance matrix Σ, which is also estimated from the data. This error must satisfy some mild statistical assumptions. The covariance matrix Σ captures the contemporaneous dependence, whereas the coefficient matrices capture the dynamic linkages present in the system. The choice of the lag length p is an empirical issue that can be selected by, e.g., the Akaike Information Criterion and ensuring that the residuals are not autocorrelated. The model is easily estimated by ordinary least squares. Based on the estimated VAR one can compute the generalized forecast error variance decomposition as follows. This requires the Vector Moving Average representation of the VAR model:

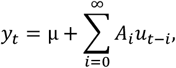

The moving average coefficients *A*_*i*_, which can straightforwardly be computed from the VAR coefficients Φ_*i*_, capture how shocks propagate through the system allowing one to understand its implied dynamics and computing interpretable measures. The one we consider here measures how much of the *h*-step forecast error variance for predicting *y*_*i*_can be attributed to shocks to *y*_*j*_. This is called the variance decomposition and a generalized version that avoids identifying structural shocks was proposed by Pesaran and Shin [31] and is formally defined as

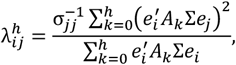

where σ_*jj*_is the standard deviation and the error for the j-th equation and *e*_*i*_is a selection vector with i-th element one and zeros everywhere else. Thus, this share depends both on the contemporaneous covariance Σ and the dynamic dependence in *A*, and therefore indirectly on Φ_*i*_. These are then normalized as by dividing each 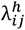 by the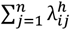, so it satisfies 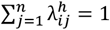 and 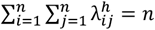, allowing us to interpret them as shares. To repeat, the (normalized) spillover measure 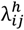 is a relative measure of variation in the variable *y*_*i*_due to shocks to the variable *y*_*j*_, where by variation we mean the forecast error variance of an *h*-period ahead prediction. These are then typically presented in a table, where the elements on the main diagonal, 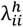, are the fraction of forecast error variance that can be explained by its own shocks to *y*_*i*_itself, whereas the remaining elements represents the fraction of the forecast error variance of the row variable due shocks to the respective column variables.

From this Diebold and Yilmaz [11] define a number of useful and interpretable spillover measures. In particular, the total spillover (TS) index is defined as the fraction of overall forecast error variance that is due to shocks to other variables, thereby measuring the degree of spillover in the system:

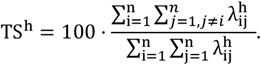

Additionally, one can define directional spillover (DS) measures, namely the spillover transmitted from variable *y*_*i*_to all other variables in the system,

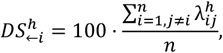

and the spillover received by variable *y*_*i*_from all other variable,

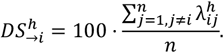

Finally, the difference between transmitted and received spillovers of *y*_*i*_defines the net spillover (NS)

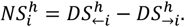

Thus, directional and net spillover measures allow assessing which brain regions are senders and which ones are receivers of information. Spillover tables then typically contain 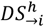 as the last column being the sum of the row elements excluding *i* and 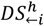 as the last row being the sum over all column elements excluding *i*. The total spillover index *TS*^*h*^ is usually presented in the bottom right corner.

In our analysis, we compute the spillover tables as defined above for resting state and working state measurements to understand how information is processed within and between regions in the brain. By contrasting working state and resting state spillover tables we aim to identify relevant and significant differences between the two in order to understand how specific memory tasks alter information transmissions. After computing the spillover measures for all 39 participants in our sample we are able to estimate the reliability of the measures themselves and also tested if spillover differences between resting and working states reached statistical significance. Due to the multiple testing nature of this problem resulting from the large number of connections, we apply a strict Bonferroni correction and graphically display those connections and changes thereof that passed those tests.

### Fit statistics of VAR models and residual autocorrelations

We estimated Akaike’s and Bayes information criterion using the DY Index MATLAB toolbox (https://github.com/binhpham79/DYIndex) and discovered that the theoretically proposed lag-1 model indeed fits the data best (**Supplementary Fig. 19**). As a last preparatory step, we conducted an analysis to determine if the residuals of VAR models were affected by serial auto correlations, as the time series for the spillover analyses were not corrected for these correlations. Our findings indicated that there were no residual autocorrelations detected (**Supplementary Fig. 20**) meaning that VAR(1) models can model the dynamics in the data sufficiently well.

### Implementation of Spillover Analysis and Filtering Procedure

For each participating subject, we imported 34 time series of the working and resting state conditions into the DY Index MATLAB toolbox (https://github.com/binhpham79/DYIndex) that was programmed by Binh Pham. We conducted a spillover analysis on the entire dataset, employing a window size of 10 time points (*h=10*), each with a repetition time (TR) of 1.24 seconds, matching the duration of the task phase (12.4 seconds). Spillover matrices were generated for all lags ranging from 1 to 5. This yielded a total of 1170 spillover matrices, derived from 3 experimental conditions, 5 lags, and 39 subjects and 2 runs. Given that the accuracy of the spillover estimates depends on the number of parameters, we applied a stringent filtering procedure to our rather large connectome consisting of 1156 relationships. A two-sided paired t-test was conducted to establish the significance of contrasts between working and resting states at the second level for each connection across all lags (1 to 5). We executed this process separately for spatial and verbal working memory contrasts. A spillover index difference was selected when the Bonferroni-corrected conjunction analysis reached significance and the intraclass correlation coefficient (ICC(2,1)) exceeded a value of 0.4 (fair reliability) in at least one of the lags. For the Bonferroni corrected conjunction analysis, we selected the largest p-value of the test and retest run, which was then subjected to Bonferroni correction at a threshold of 0.05 divided by the number of spillover analyses (1156) assuming a two-sided test (0.05/(1156*2)). By employing this filter procedure, we reduced the connectome from the original 1156 potential connections down to 196 (from a 34×34 to a 14×14 matrix). Subsequently, we recalculated the spillover coefficients for this refined connectome, along with their test-retest reliability estimates for lags 1 through 5 which again resulted in 1170 spillover matrices. We contrasted the working state spillover maps with the resting state maps and selected a spillover when the test and retest run reached a Bonferroni corrected p value (conjunction analysis). In addition, the difference between working and resting state spillover should exhibit at least a test-retest reliability of ICC(2,1) >0.4. As NHST makes little sense in spillover research we also selected a path when a mean spillover of 5 was reached with a test-retest reliability of ICC> 0.6. It should be mentioned that the test-retest reliability of narrow contrasts is indeed substantially lower when compared to the experimental conditions that are not subjected to narrow contrast.

### Test-Retest Reliability Assessment

A larger family of intra class correlation coefficients that can be used to estimate the reliability of empirical data [32]. We utilized a two-way random, single score model to quantify the path-specific test-retest reliability of conventional connectivity and spillover metrics which is advised for imaging purposes [18]. The 39 observations of the test run were correlated with 39 observations of the retest run, yielding 561 correlations for classic metrics and 1156 for spillover data. Following Cicchetti’s criteria, ICC values were categorized as follows: less than 0.40 indicates poor reliability, 0.40 to 0.59 suggests fair, 0.60 to 0.74 shows good, and greater than 0.75 represents excellent reliability [22]. We ascertained the reproducibility of these measures separately for each of the 3 experimental conditions and for the contrast between the working and resting states by entering their respective differences into the ICC model.

### Consistency of Spillover Maps Across Time Lags

Establishing the homogeneity of spillover maps generated by various lag models was essential. To determine consistency, we averaged maps across individuals per run per lag. The two-dimensional spillover matrices were transferred into one dimension. The resulting spill vectors were correlated across the different lags per experimental condition. The outlying nature of within regions spillover may artificially inflate cross-lag correlations. To account for this we included all spillovers in addition to a version excluding intra-regional spillover, labeled “No Diag.”, and we report these findings in **Supplementary Table 1**. We further assessed the correlation consistency for working minus resting state contrasts across different lags, as documented in **Supplementary Table 1**. These results reveal a high degree of similarity among maps, notwithstanding a marginal decline in map likeness concomitant with increased lags. Scatterplots of these contrasts across various lags are presented in **Supplementary Fig. 2-7**, illustrating the nominal decrease in similarity possibly due to the elevated complexity and estimation error attributable to the larger number of parameters at higher lags.

### Convergence of Working and Resting State Spillover Maps at Lag 1

Given the high across lag stability of spillover maps we decided to limit this analysis to lag 1 spillover only. We averaged spillover maps for each run and scattered these averages across experimental conditions. Distinct scatterplots were generated for spillover values between regions (**Supplementary Fig. 15**) versus those within regions (**Supplementary Fig. 16**) to illustrate the persistent robustness despite the extreme values of intra-regional spillovers. We proceeded to compute the average of test and retest maps and performed regression analysis across the three experimental conditions under study, as outlined in **Supplementary Fig. 14**.

### Correlation Between Conventional Connectivity and Spillover Maps

To facilitate the interpretation of spillover maps, we compared averaged conventional connectivity maps with averaged spillover maps across experimental conditions. It is important to note that the conventional connectivity maps signify unidirectional relationships, whereas spillover maps represent bidirectional interactions. Additionally, while conventional maps inherently carry a correlation of one within a brain region, spillover maps do not. We addressed this discrepancy by correlating the lower triangular part of the connectivity matrix with the lower and upper triangular part of the spillover matrix that are related to sending and receiving behavior. Resultant scatterplots are provided in **Supplementary Fig. 17** (sending) and **Supplementary Fig. 18** (receiving) with correlation coefficients detailed in **Supplementary Table 4**.

### Cognitive load and Resting load

It reasonable to assume that the neurocognitive load of particular experiential condition is related to the amount of information that is circulated between brain regions. The latter can be described in terms of total spillover TS^h^ which captures the amount of information that is processed among brain regions relative to the total information processed. We tested whether working and resting state conditions where significantly different from each other by means of conjunction analysis and also estimated test-retest reliability ICC(2,1).

In a next step we estimated the relative increase in cognitive load when working state was compared to resting state conditions:

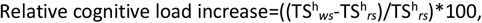

where TS^h^ denotes the total spillover and *ws* and *rs* denote working and resting state respectively.

As previously discussed, we did not test spillover statistics of the resting brain against zero by means of null significance hypothesis testing as this equals the hypothesis that the resting brain processes more information than the dead brain which is most likely the case. However, it might be interesting to assess how the resting brain state compares to working state conditions and dead brain conditions. As mentioned, we make the not unlikely assumption that the dead brain processes zero information. We assume that the total information processed among brain regions during resting state in comparison to the dead brain equals the amount of neural information processed during rest. Furthermore, we assume that the amount of information of the working brain in comparison to the resting brain equals the relative amount of neural information that is needed to execute a cognitive task. We will refer to this difference as *neurocognitive load*. Given that the neuro cognitive load is known it is possible to estimate how much neural information is processed during resting state. We will refer to this as resting load:

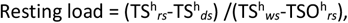

where *ds* denotes the dead brain state which is assumed to be zero (*ds*=0).

## Ethics approval

All individuals provided written informed consent. The research was executed according to the guidelines of the Declaration of Helsinki. The study was approved by the Ethics committee under GZ 39/31/63 ex 2011/12.

## Data availability statement

The anonymized data collected are available as matlab files via the github data repository at: https://github.com/hinata2305/SpillOver

## References

1. Cowan, N. What are the differences between long-term, short-term, and working memory? Progress in brain research 169, 323–338 (2008).

2. Diamond, A. Executive functions. Annual review of psychology 64, 135–168 (2013).

3. Postle, B. R. Working memory as an emergent property of the mind and brain. Neuroscience 139, 23–38 (2006).

4. Miller, E. K., Lundqvist, M. & Bastos, A. M. Working Memory 2.0. Neuron 100, 463–475 (2018).

5. Bays, P. M., Schneegans, S., Ma, W. J. & Brady, T. F. Representation and computation in visual working memory. Nature human behaviour 8, 1016–1034 (2024).

6. Ramsey, J. D. et al./person-group>. Six problems for causal inference from fMRI. NeuroImage 49, 1545–1558 (2010).

7. Parente, F. & Colosimo, A. Modelling a multiplex brain network by local transfer entropy. Scientific reports 11 (2021).

8. Mynbaev, K. T. Short-memory linear processes and econometric applications (Wiley, 2011).

9. Beran, J. Statistics for Long-Memory Processes (Routledge, 2017).

10. Diebold, F. X. & Yilmaz, K. Measuring Financial Asset Return and Volatility Spillovers, with Application to Global Equity Markets. The Economic Journal 119, 158–171 (2009).

11. Diebold, F. X. & Yilmaz, K. Better to give than to receive: Predictive directional measurement of volatility spillovers. International Journal of Forecasting 28, 57–66 (2012).

12. Baddeley, A. D. Is working memory still working? The American psychologist 56, 851–864 (2001).

13. Coppola, P. et al./person-group>. The complexity of the stream of consciousness. Communications Biology 5, 1173 (2022).

14. James, W. The Principles of Psychology (Henry Holt, 1890).

15. Costall, A. ‘Introspectionism’ and the mythical origins of scientific psychology. Consciousness and cognition 15, 634–654 (2006).

16. Nee, D. E. et al./person-group>. A meta-analysis of executive components of working memory. Cerebral cortex (New York, N.Y. : 1991) 23, 264–282 (2013).

17. Noble, S., Scheinost, D. & Constable, R. T. A decade of test-retest reliability of functional connectivity: A systematic review and meta-analysis. NeuroImage 203, 116157 (2019).

18. Noble, S., Scheinost, D. & Constable, R. T. A guide to the measurement and interpretation of fMRI test-retest reliability. Current opinion in behavioral sciences 40, 27–32 (2021).

19. Elliott, M. L. et al./person-group>. What Is the Test-Retest Reliability of Common Task-Functional MRI Measures? New Empirical Evidence and a Meta-Analysis. Psychological science 31, 792–806 (2020).

20. Dinkel, P. J., Willmes, K., Krinzinger, H., Konrad, K. & Koten, J. W. Diagnosing developmental dyscalculia on the basis of reliable single case FMRI methods: promises and limitations. PloS one 8, e83722 (2013).

21. Friston, K. J., Penny, W. D. & Glaser, D. E. Conjunction revisited. NeuroImage 25, 661–667 (2005).

22. Cicchetti, D. V. Guidelines, criteria, and rules of thumb for evaluating normed and standardized assessment instruments in psychology. Psychological Assessment 6, 284–290 (1994).

23. Funder, D. C. & Ozer, D. J. Evaluating Effect Size in Psychological Research: Sense and Nonsense. Advances in Methods and Practices in Psychological Science 2, 156–168 (2019).

24. Nolan, L. (ed.). The Cambridge Descartes Lexicon (Cambridge University Press, 2016).

25. in Historical and Philosophical Foundations of Psychology, edited by M. Farrell (Cambridge University Press, 2018), pp. 70–93.

26. FreeSurfer. Harvard. Available at https://surfer.nmr.mgh.harvard.edu/.

27. CleanBrain. Koten, Jan Willem; Schüppen, Andre. Available at https://github.com/hinata2305/CleanBrain.

28. Alakörkkö, T., Saarimäki, H., Glerean, E., Saramäki, J. & Korhonen, O. Effects of spatial smoothing on functional brain networks. The European Journal of Neuroscience 46, 2471–2480 (2017).

29. Hagler, D. J., Saygin, A. P. & Sereno, M. I. Smoothing and cluster thresholding for cortical surface-based group analysis of fMRI data. NeuroImage 33, 1093–1103 (2006).

30. Olszowy, W., Aston, J., Rua, C. & Williams, G. B. Accurate autocorrelation modeling substantially improves fMRI reliability. Nature communications 10, 1220 (2019).

31. Pesaran, H. & Shin, Y. Generalized impulse response analysis in linear multivariate models. Economics Letters 58, 17–29 (1998).

32. Shrout, P. E. & Fleiss, J. L. Intraclass correlations: uses in assessing rater reliability. Psychological bulletin 86, 420–428 (1979).

