## Supplementary Material for "Information Spillover in ‘Resting Memory’ and ‘Working Memory’"

**Supplementary Figure 1: Reliability of conventional connectivity maps**

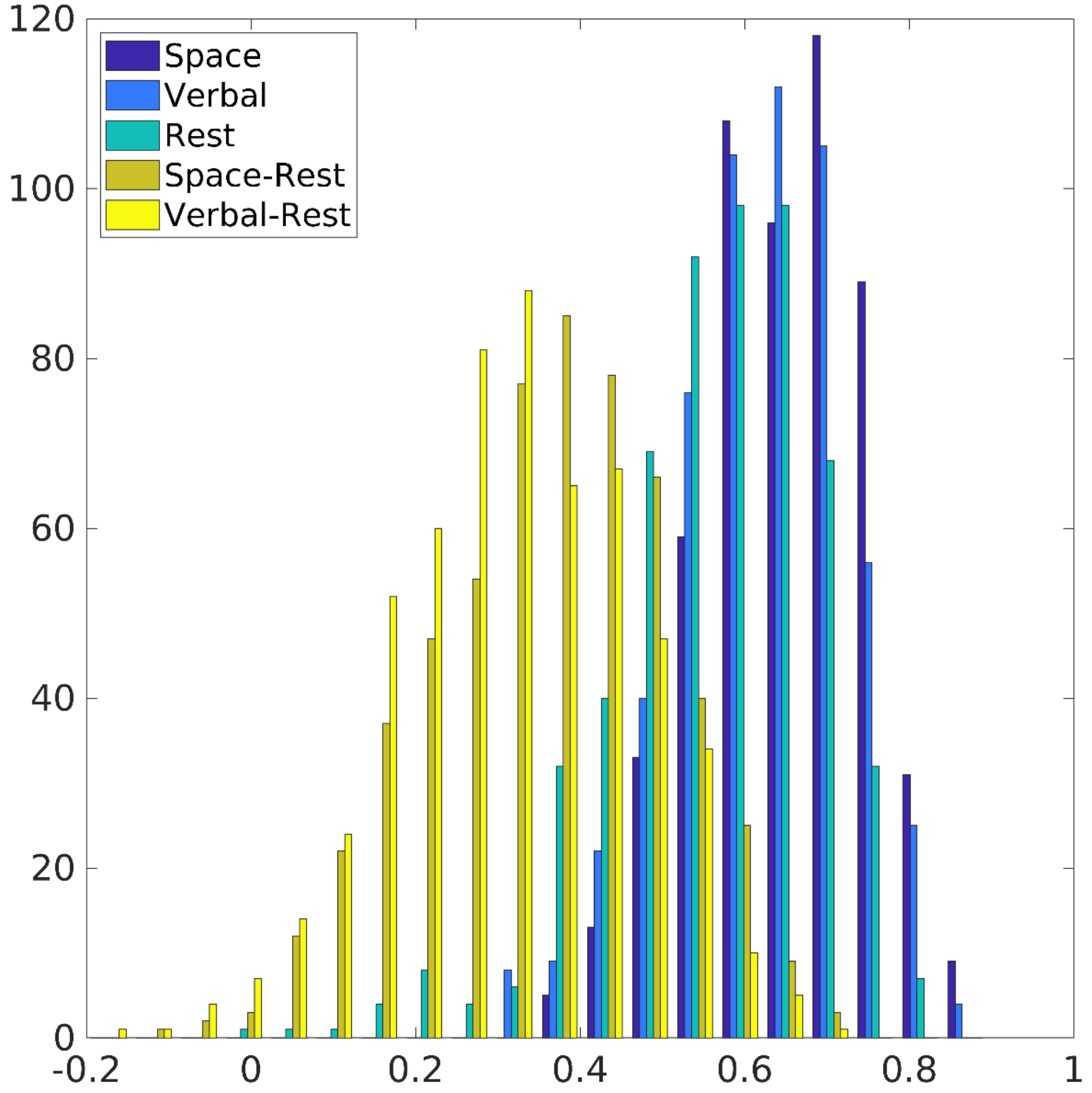


**A path wise test-retest reliability analysis was estimated employing an ICC(2,1) model. This model estimates whether differences in connectivity strength between individuals remain conserved over time. The mean test-retest reliability of the 561 paths under study were 0.66 for spatial WM; 0.64 for the verbal WM and 0.58 for the resting state data. However, the mean connectome reliability dropped to 0.36 and 0.32 when spatial working memory and verbal working memory were contrasted with resting state data respectively. For this analysis resting state connectivity was subtracted from working state connectivity before correlations were entered into the ICC(2,1) model. Note that Fisher z-transformations were applied whenever a path correlation was subjected to statical analysis.**

#### Supplementary Figure 2: Across lag scatter plots of the contrast Spatial WM minus Rest for Test run

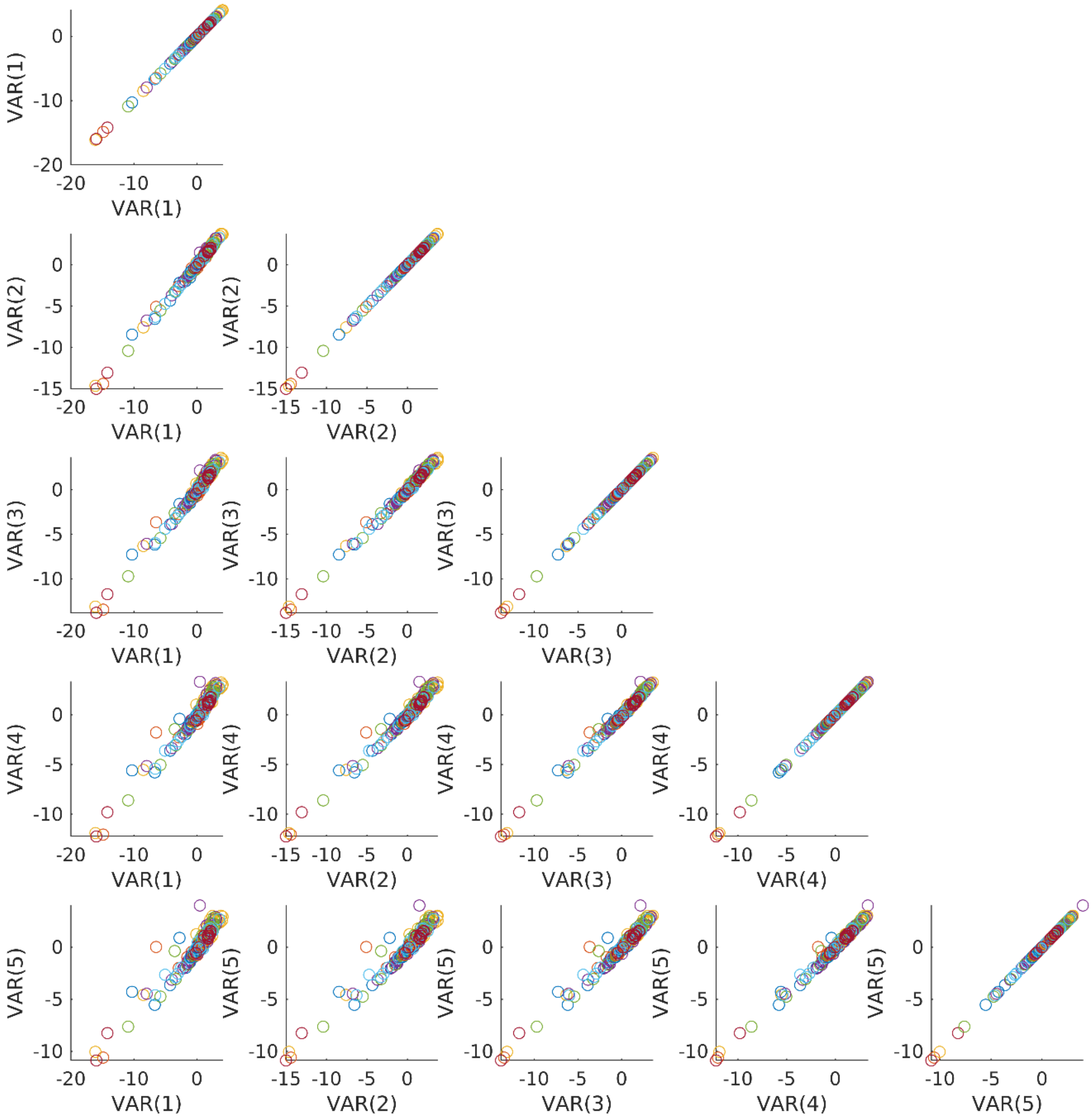


**Spillover maps were estimated for VAR(1) till VAR(5). We averaged the spillover maps across individuals per experimental condition per run per lag. Subsequently we subtracted the resting state spillover maps from the spatial working memory spillover maps obtained from the test run per lag. The 196 spillover statistics per map were scattered across lags. The horizontal and vertical axis report the VAR models that were scattered.**

**Supplementary Figure 3: Across lag scatter plots of the contrast Spatial WM-minus-Rest for Retest run**

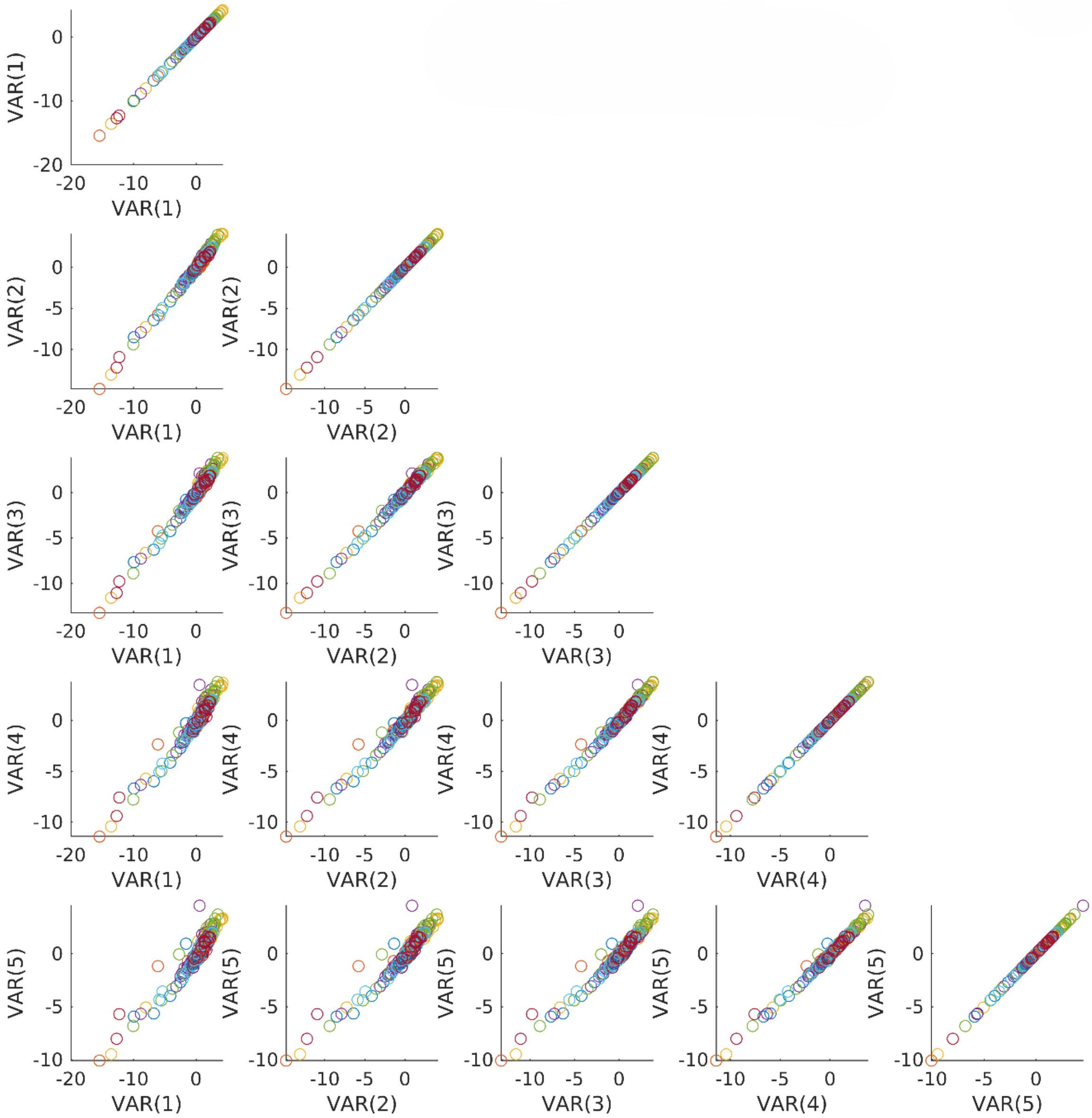


**Spillover maps were estimated for VAR(1) till VAR(5). We averaged the spillover maps across individuals per experimental condition per run per lag. Subsequently we subtracted the resting state spillover maps from the spatial working memory spillover maps obtained from the retest run per lag. The 196 spillover statistics per map were scattered across lags. The horizontal and vertical axis report the VAR models that were scattered.**

#### Supplementary Figure 4: Across lag scatter plots of the contrast Verbal WM-minus-Rest for Test run

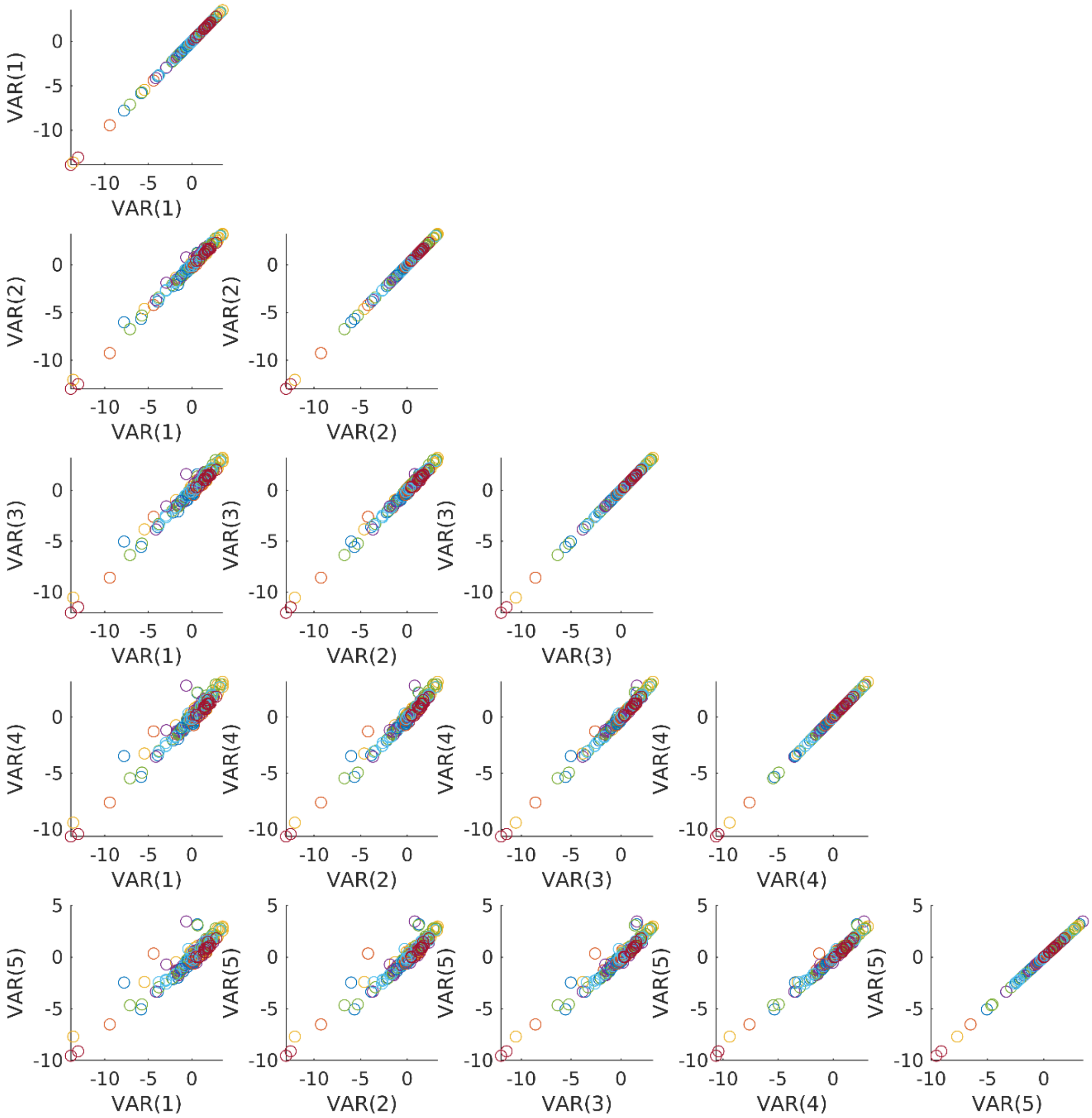


**Spillover maps were estimated for VAR(1) till VAR(5). We averaged the spillover maps across individuals per experimental condition per run per lag. Subsequently we subtracted the resting state spillover maps from the verbal working memory spillover maps obtained from the test run per lag. The 196 spillover statistics per map were scattered across lags. The horizontal and vertical axis report the VAR models that were scattered.**

#### Supplementary Figure 5: Across lag scatter plots of the contrast Verbal WM-minus-Rest for Retest run

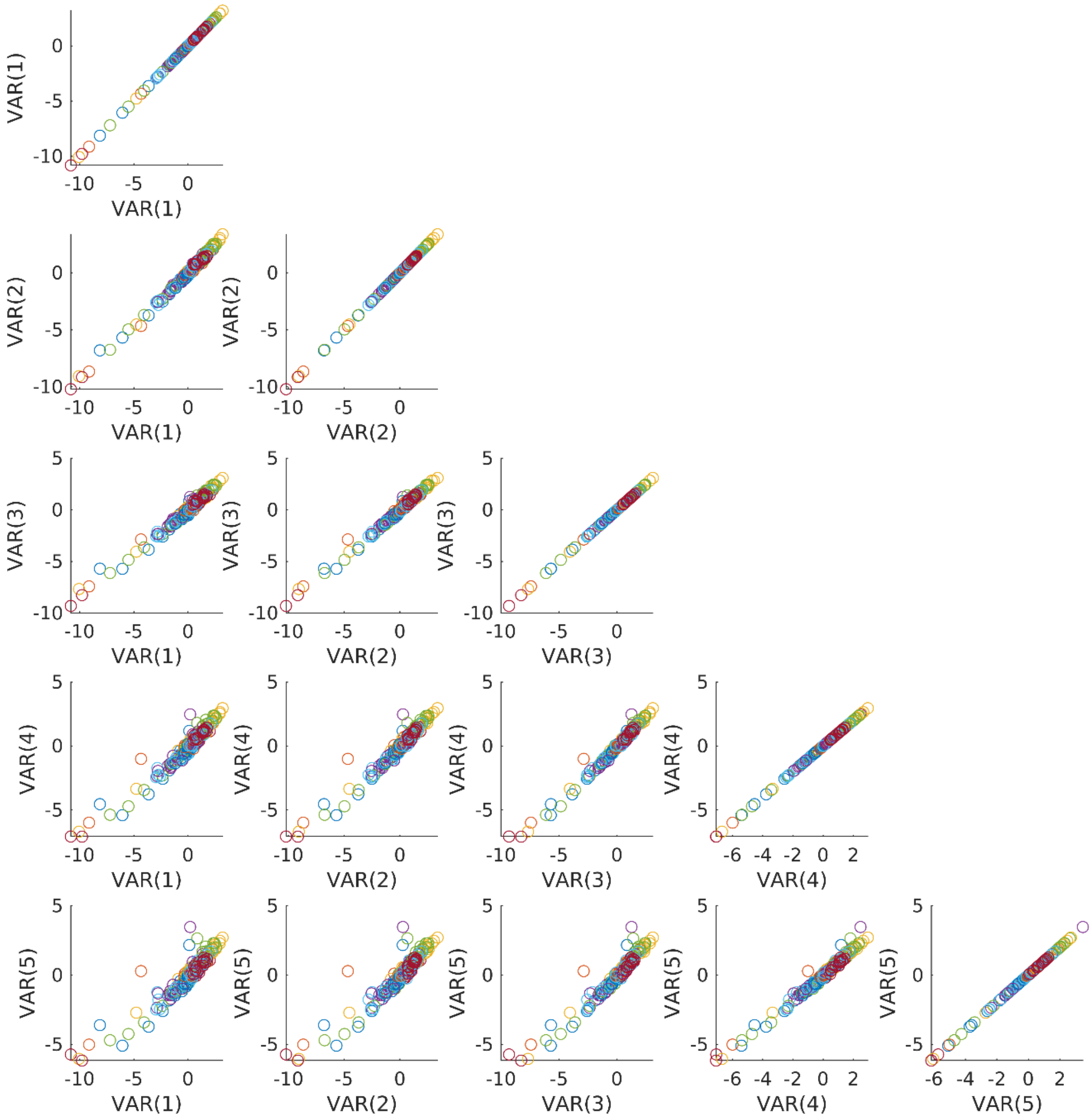


**Spillover maps were estimated for VAR(1) till VAR(5). We averaged the spillover maps across individuals per experimental condition per run per lag. Subsequently we subtracted the resting state spillover maps from the verbal working memory spillover maps obtained from the retest run per lag. The 196 spillover statistics per map were scattered across lags. The horizontal and vertical axis report the VAR models that were scattered.**

#### Supplementary Figure 6: Across lag scatter plots of the contrast Spatial WM-minus-Verbal WM for Test run

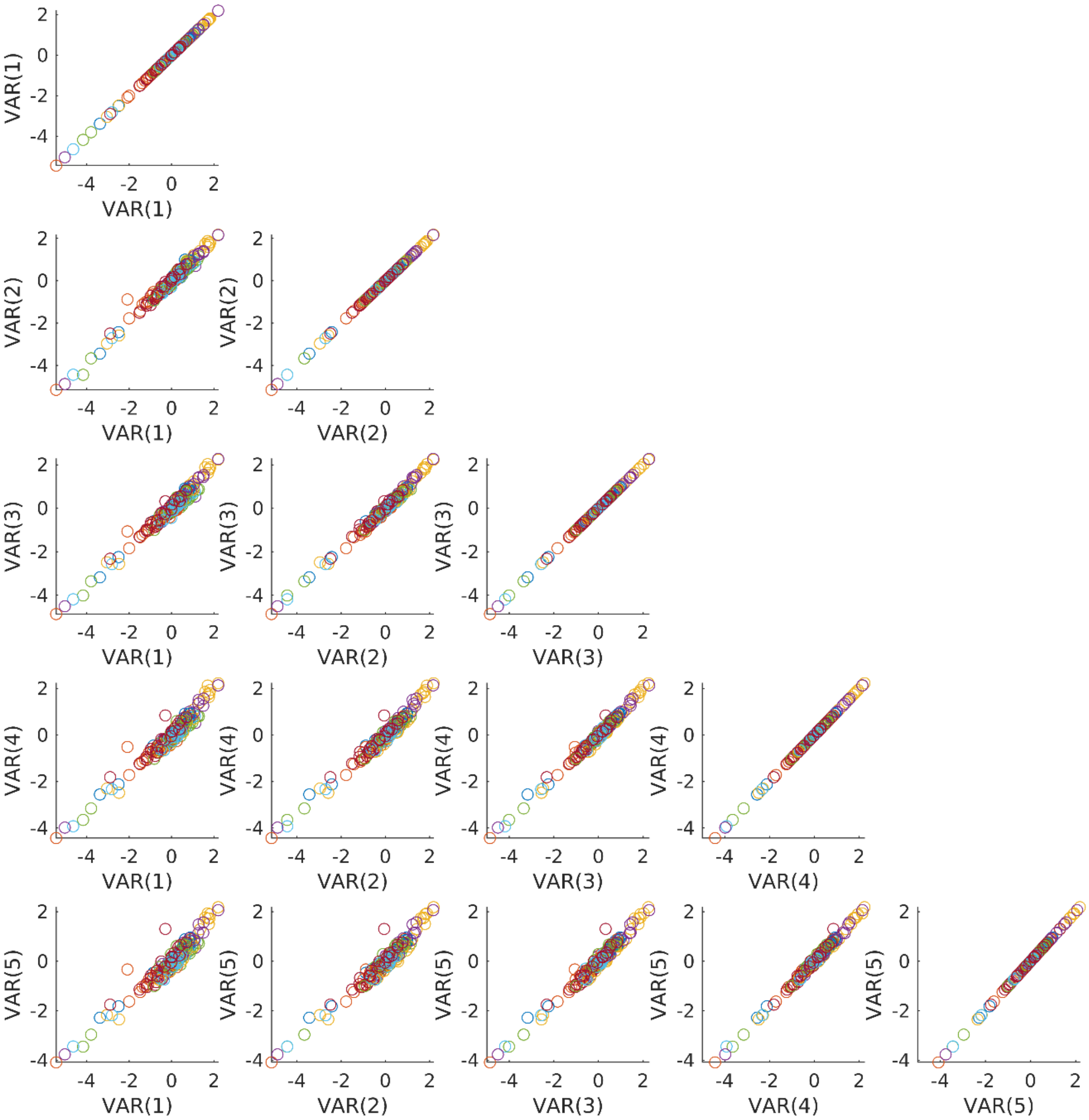


**Spillover maps were estimated for VAR(1) till VAR(5). We averaged the spillover maps across individuals per experimental condition per run per lag. Subsequently we subtracted the spatial spillover maps from the verbal spillover maps obtained from the test run per lag. The 196 spillover statistics per map were scattered across lags. The horizontal and vertical axis report the VAR models that were scattered.**

**Supplementary Figure 7: Across lag scatter plots of the contrast Spatial WM-minus-Verbal WM for Retest run**

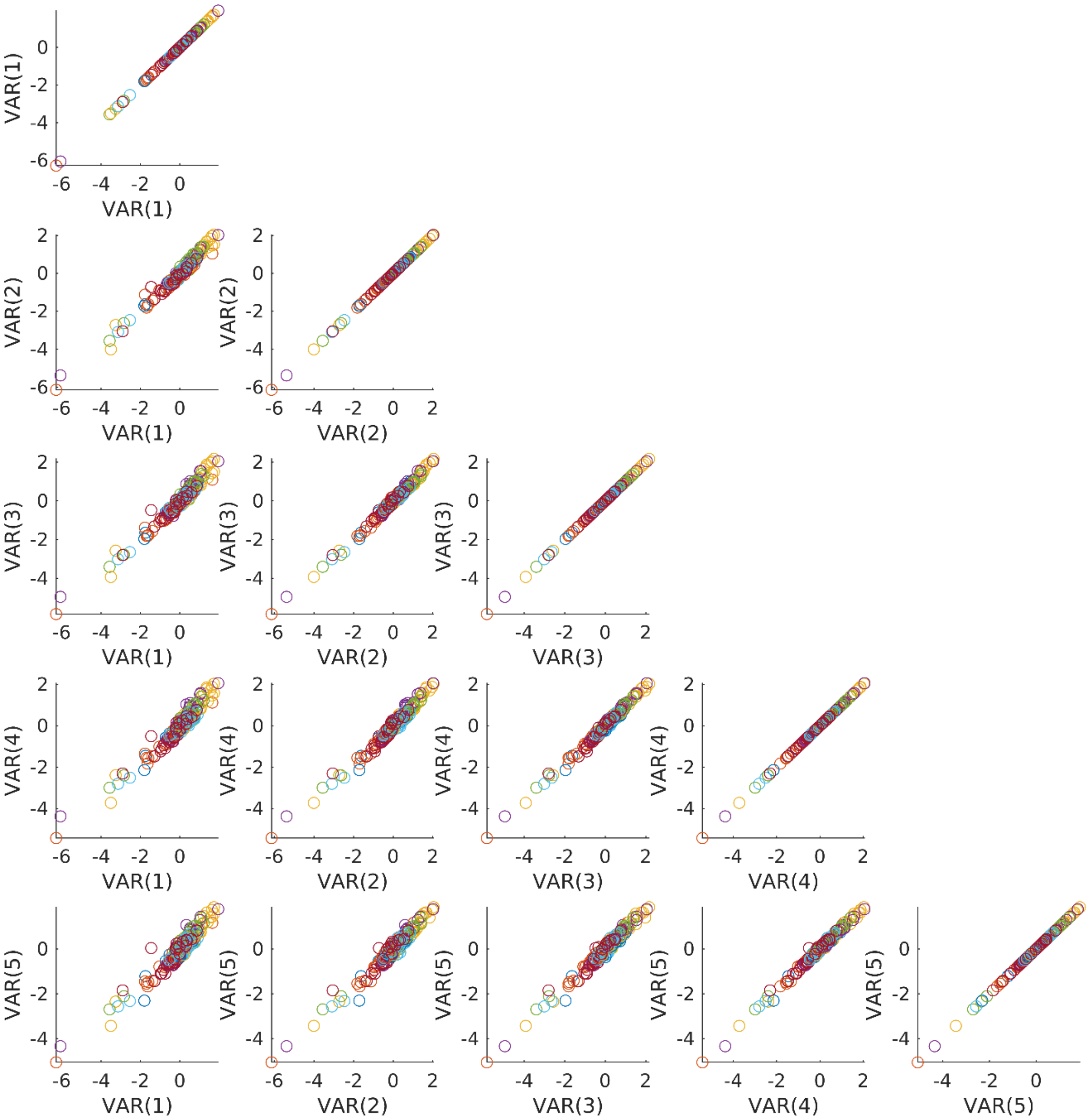


**Spillover maps were estimated for VAR(1) till VAR(5). We averaged the spillover maps across individuals per experimental condition per run per lag. Subsequently we subtracted the spatial spillover maps from the verbal spillover maps obtained from the retest run per lag. The 196 spillover statistics per map were scattered across lags. The horizontal and vertical axis report the VAR models that were scattered.**

#### Supplementary Figure 8: Task related signal averages

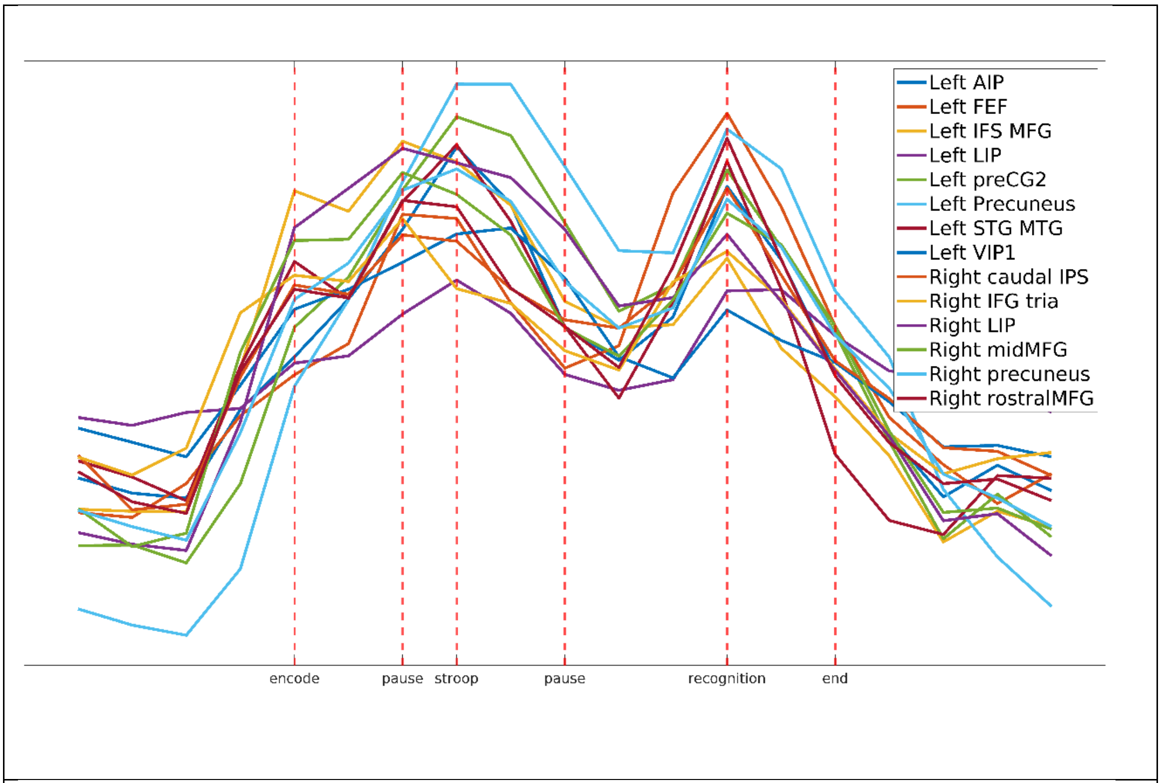


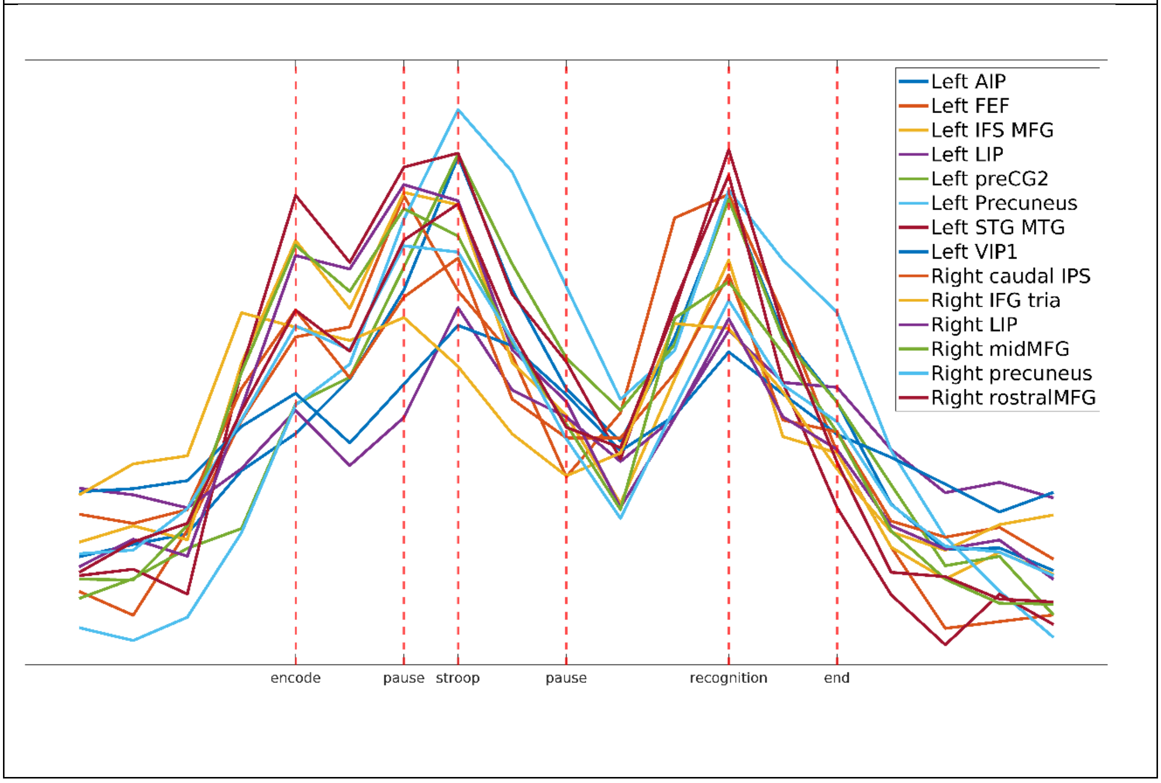


**The 24 working memory related signals of an experiment were averaged across the sample and runs per brain region. The task related signal averages of the 14 time courses of interest are depicted per task. Top the spatial WM task bottom the verbal WM task. Assuming a time to peak of 6 seconds we were able link the cognitive component to the time course. Cognitive components are reported on the horizonal axis.**

**Supplementary Figure 9 : Comparing task related signal averages in test and retest runs to the overall brain response during a spatial WM task.**

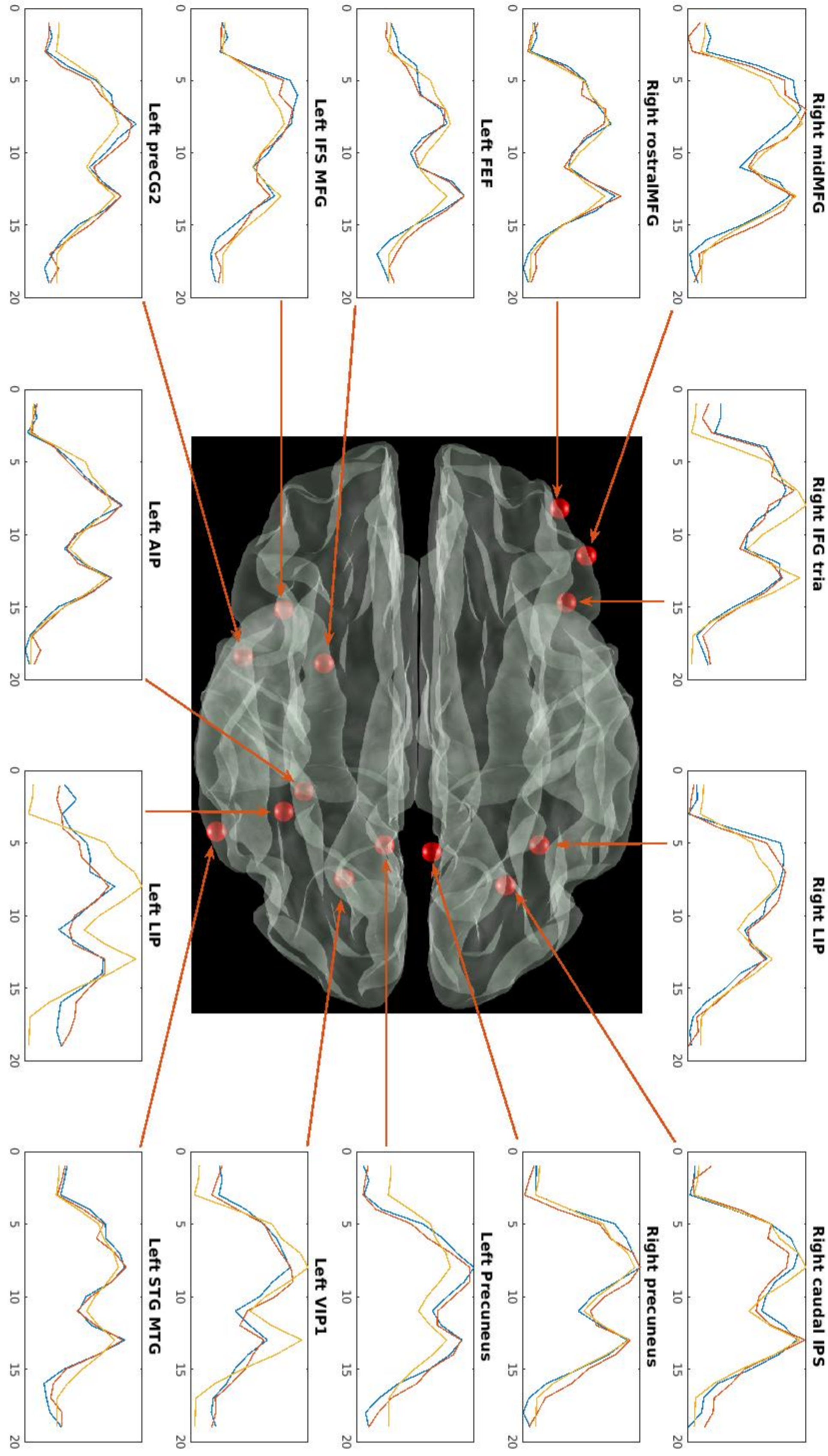


**This figure shows task related signal averages of the spatial WM task. It illustrates the overall grand mean task related signal average of 14 regions from a test and retest run, shown in yellow. Additionally, the task related averages from the test and retest runs were separately estimated for each region and depicted in red and blue, respectively. More details can be found in the extended caption to Supplementary Figure 9 and Supplementary Figure 10.**

**Supplementary Figure 10: Comparing task related signal averages in test and retest runs to the overall brain response during a verbal WM task.**

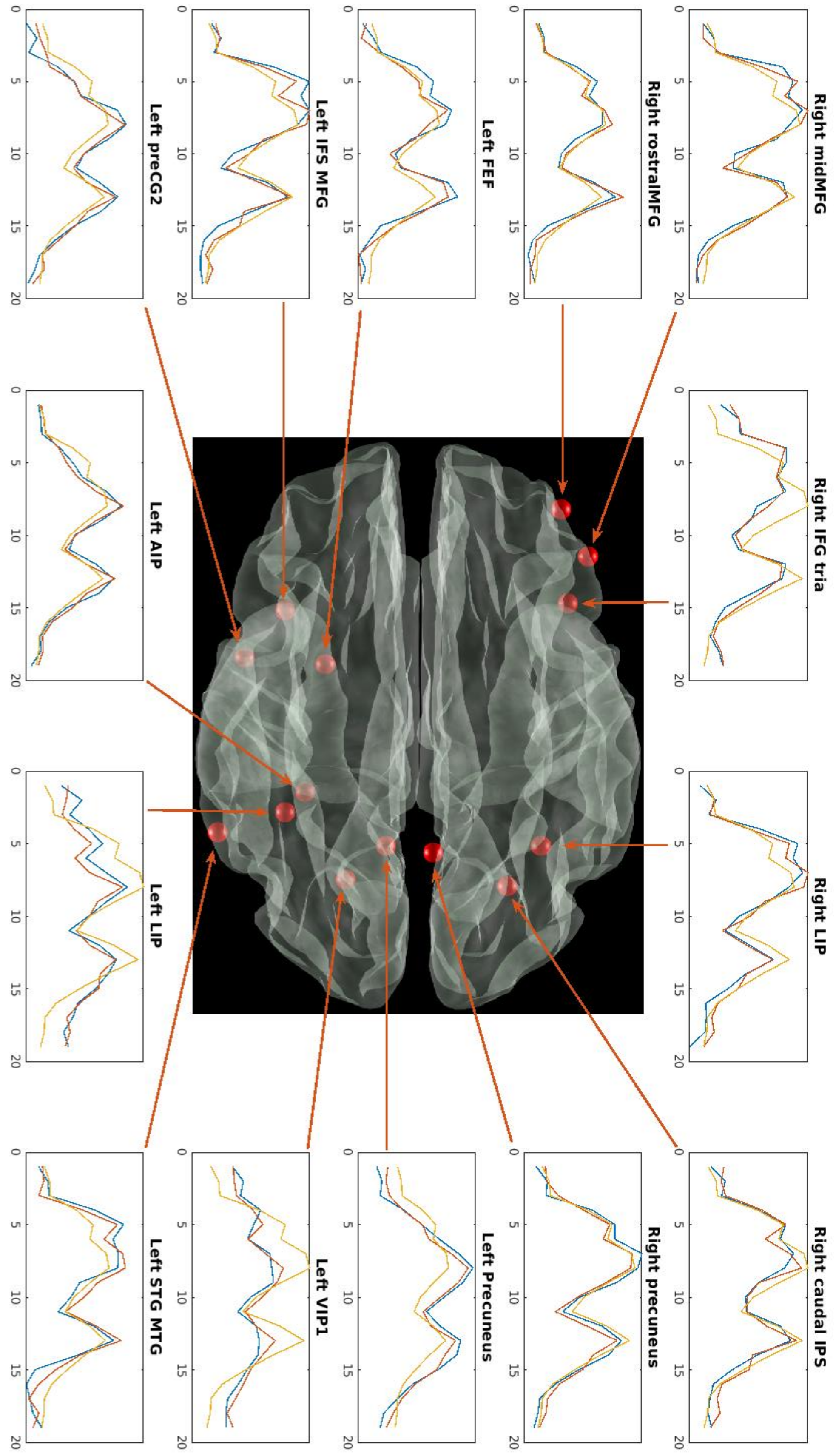


**This figure shows task related signal averages of the verbal WM task. It illustrates the overall grand mean task related signal average of 14 regions from a test and retest run, shown in yellow. Additionally, the task related averages from the test and retest runs were separately estimated for each region and depicted in red and blue, respectively. More details can be found in the extended caption to Supplementary Figure 9 and Supplementary Figure 10.**

#### Extended caption to Supplementary Figure 9 and Supplementary Figure 10

**Supplementary Figure 9 and Supplementary Figure 10 reveal that substantial differences in average time course behavior exist, suggesting that the data are potentially suitable for VAR analysis. The right inferior frontal gyrus is a rapid responder in the sense that it reaches its brain activity peak before all the other regions as can be seen from the figures. It is not unlikely that bold expression of this area is triggered by the presentation of the encoding stimuli that warns the participant that the experiment is commencing. Similarly, the left inferior frontal sulcus/middle frontal gyrus and the left lateral aspect of the IPS find their highest peaks during the early encoding phase of the experiment. As the left inferior frontal sulcus/middle frontal gyrus is roughly equivalent with left dorsolateral prefrontal cortex it is deemed important for working memory. However, the DLPFC did not necessarily exhibit sustained brain activity during the pause that followed the distraction phase. Such sustained brain activity was mostly indeed found in the left precuneus for both the verbal and spatial working memory task. It is important to notice that the left precuneus is an area that responds late to the incoming stimuli. We also observed some task specialization. While the spatial task showed some expected sustained activity in the right hemisphere more specifically the lateral aspect of IPS (Supplementary Figure 9) the opposite was true for the verbal task where increased sustained activity in the left superior temporal gyrus (Wernicke's area) and the left precentral gyrus was observed during encoding (Supplementary Figure 10). The left lateral intraparietal sulcus and the left ventral intra parietal sulcus can be characterized as later responders as they find their peaks during the performance of the number Stroop tasks that has repeatedly been shown to activates these areas. Finally, the frontal eye fields exhibit an increase in signal strength shortly before the initiation phase of the experiment commences. This suggests that participants anticipate eye movement or refresh memory traces.**

#### Supplementary Figure 11: Reliability of spillover maps

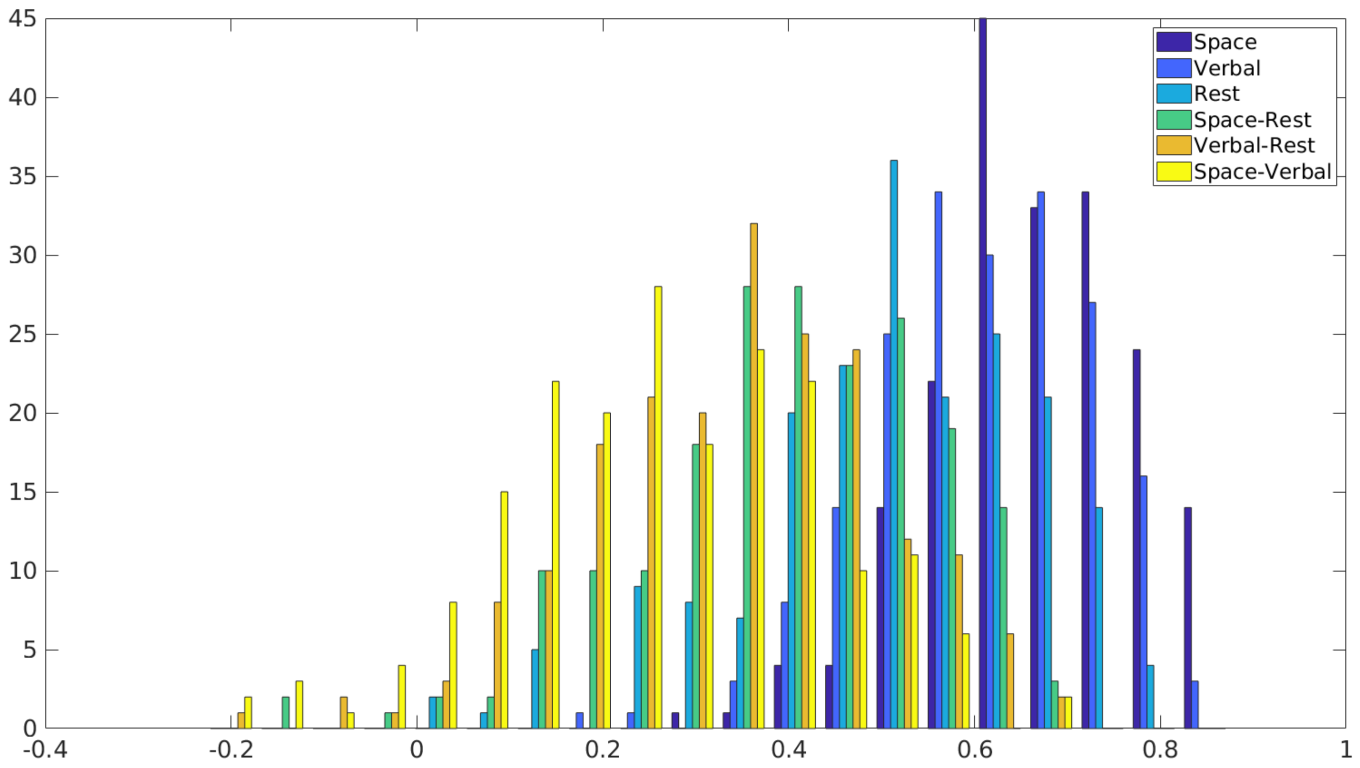


**A spillover-wise reliability analysis was estimated employing an ICC(2,1) model. This ICC model estimates whether differences in connectivity strength between individuals remain conserved over time. The pathwise test-retest reliability for the 196 spillover paths were estimated. The average test-retest reliability was 0.68 for spatial WM; 0.63 for verbal WM and 0.53 for resting state data. The mean reliability for complex contrasts was 0.41 for spatial WM-rest; 0.35 for verbal WM-rest and 0.27 for spatial WM-verbal WM.**

#### Supplementary Figure 12: Narrow Contrast

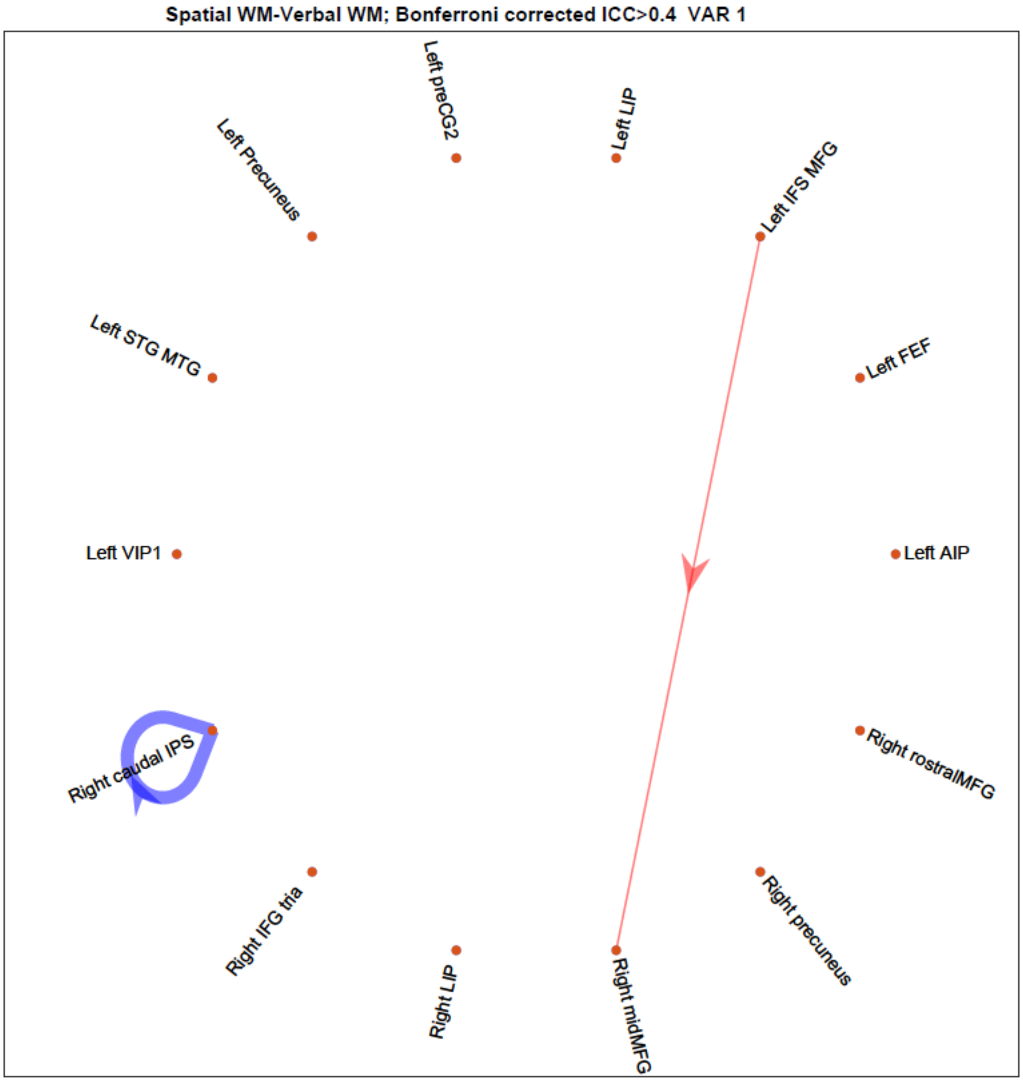


This graph provides an overview of the spillover contrast spatial WM versus verbal WM. Red arrows indicate a relative increase in spillover compared to verbal WM, while blue arrows signify a relative decrease. The differences in spillover between the states are reported in the nearby vicinity of the arrows. To be included in the results, both the initial test and subsequent retest runs had to exceed a Bonferroni-corrected threshold value, and each individual connection had to display a minimum ICC of 0.4 for the contrast in question. The results suggest that there is increased spill over from the Left IFS /MFG to the Right midMFG and also that self-referential activity in the caudal aspect of the right IPS is suppressed.

#### Supplementary Figure 13: Correlation between resting and working state in conventional connectivity

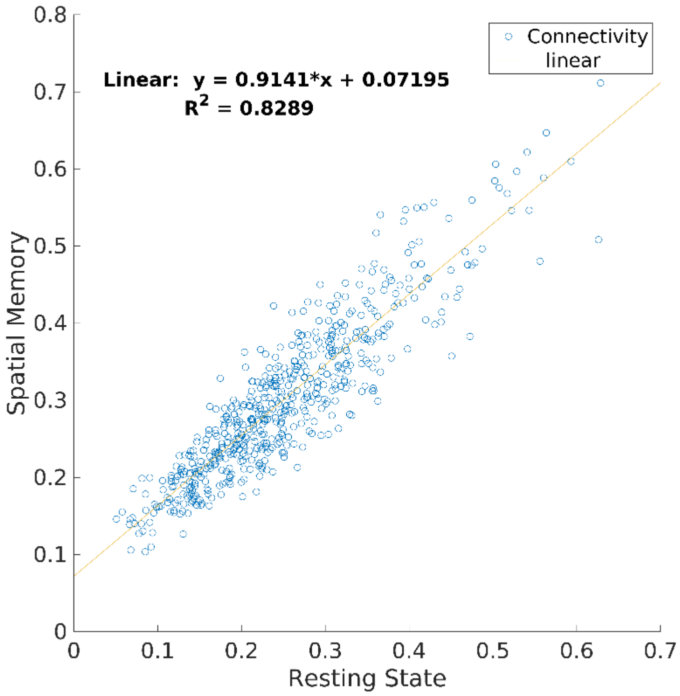


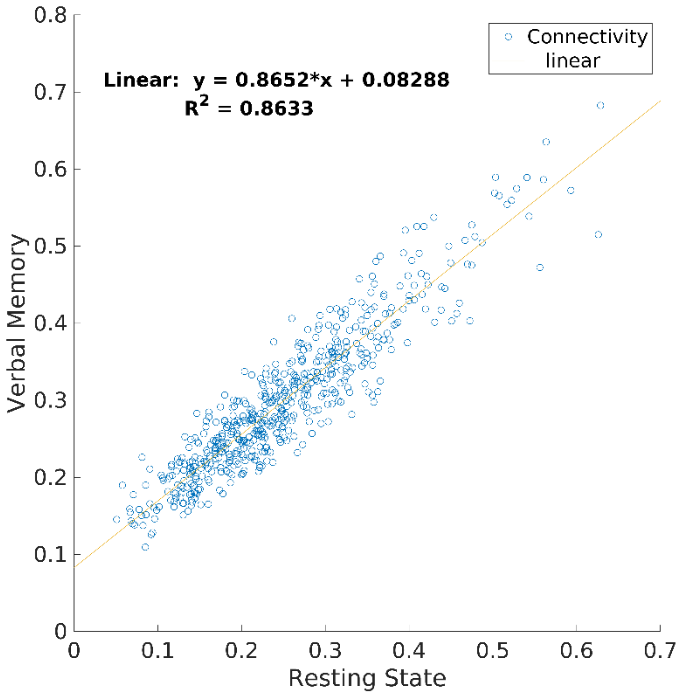


**A regression analysis conducted on the average working and resting state connectomes consisting of 561 paths showed a strong relation between the two ($R^2$=0.83 for spatial WM and $R^2$=0.86 for working WM), with intercepts of 0.072 and 0.083 respectively. This indicates that the networks involved in working state processes overlap with those involved in resting state while at the same time brain connectivity during working state is somewhat higher than during resting state.**

#### Supplementary Figure 14: Spill over scatterplots of Spatial WM, Verbal WM and resting state

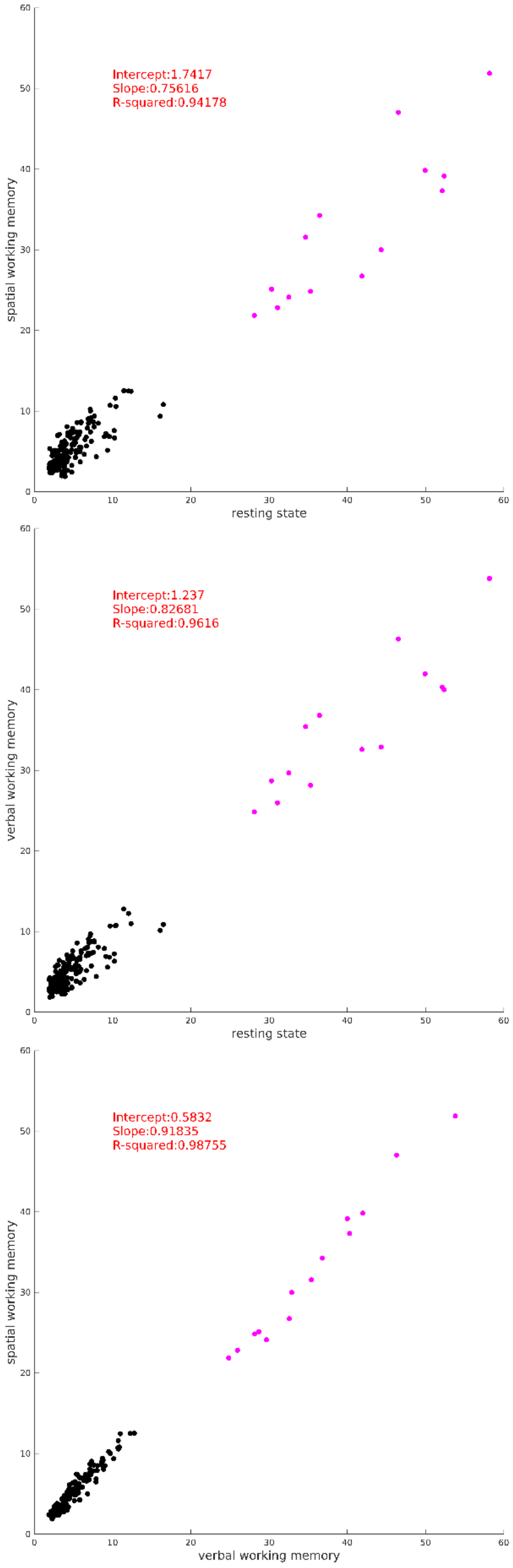


**Spillover maps were averaged across subjects and run for each task, resulting in 192 spillover values. The magenta dots represent within-region spillover, while the black dots represent between-region spillover effects. The respective tasks are given on the axis.**

#### Supplementary Figure 15: Between regions spillover scatterplots across Spatial WM, Verbal WM and resting state

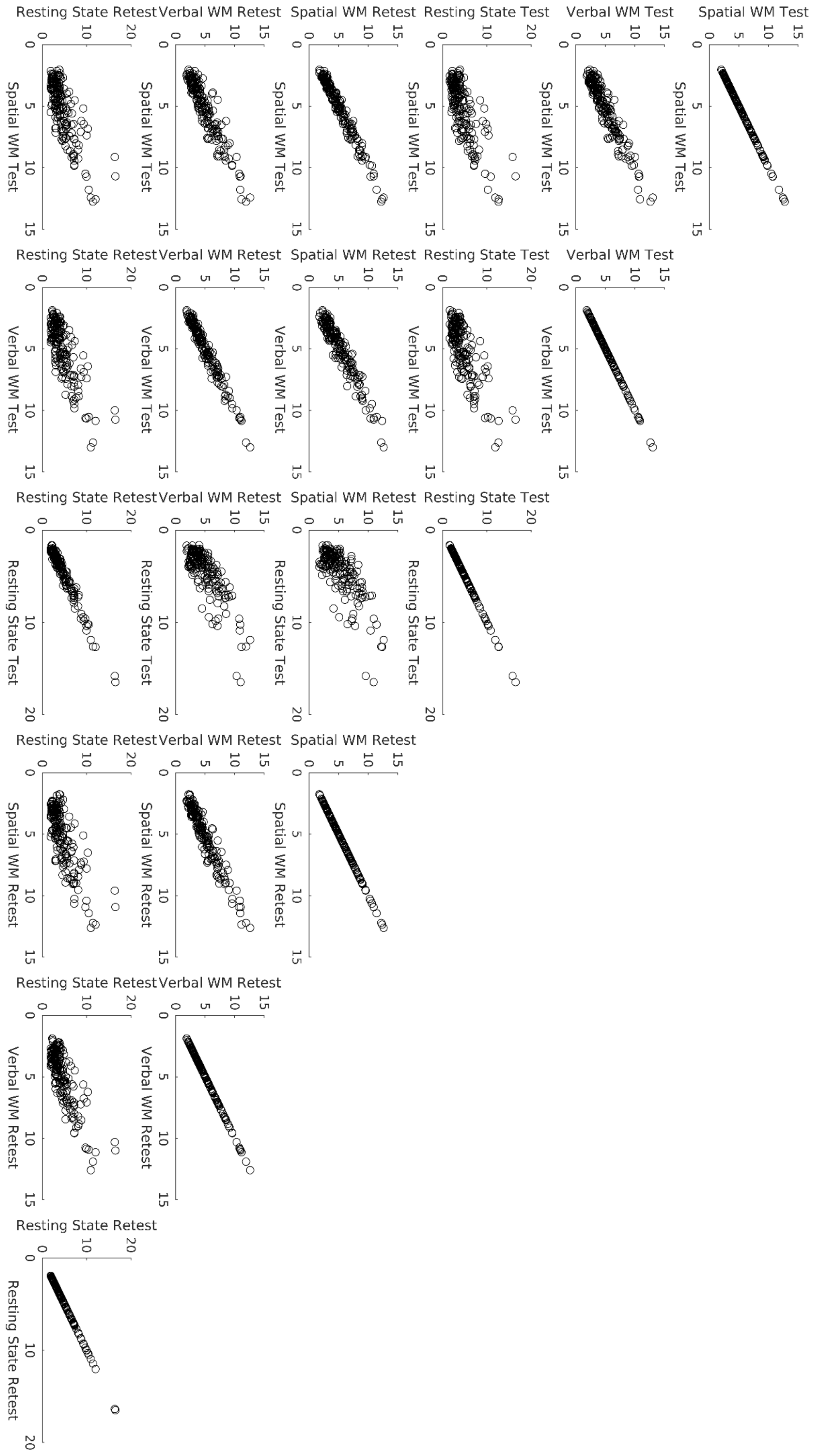


**Between region spillover maps were averaged across subjects per task per run. Subsequently, the averaged maps were scattered across tasks and runs. The tasks and runs are reported on the axis.**

#### Supplementary Figure 16: Within region spillover scatterplots across Spatial WM, Verbal WM and resting state

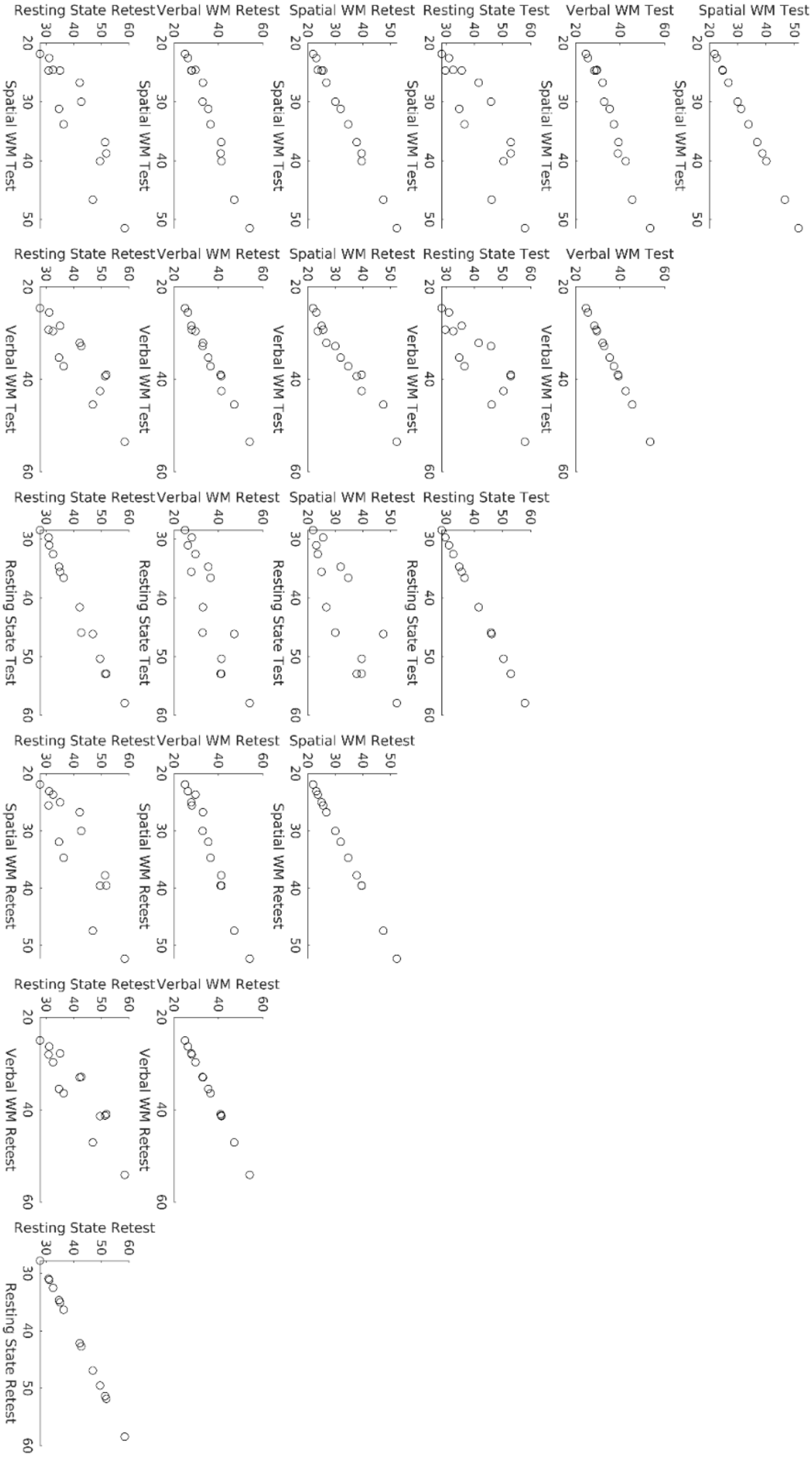


**Between region spillover maps were averaged across subjects per task per run. Subsequently, the averaged maps were scattered across tasks and runs. The tasks and runs are reported on the axis.**

#### Supplementary Figure 17: Scatter plots across sending spillover maps and connectivity maps

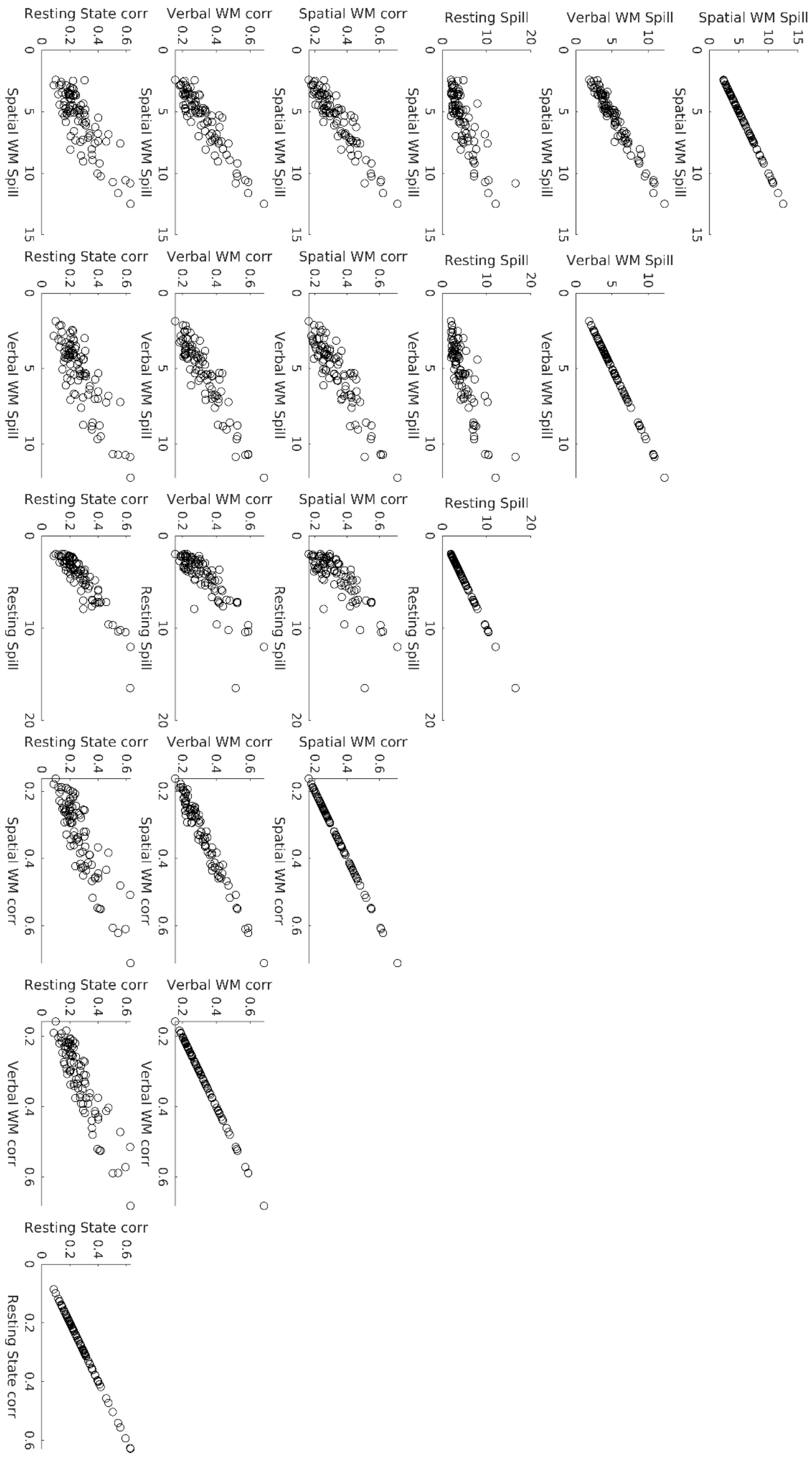

**Conventional connectivity and spillover maps were averaged across individuals. We then correlated the lower triangular sections of the connectivity maps with those of the spillover maps pertaining to the sending of information. The axes of the plots denote the task and map type (spillover or connectivity).**

#### Supplementary Figure 18: Scatter plots across receiving spill over maps and connectivity maps

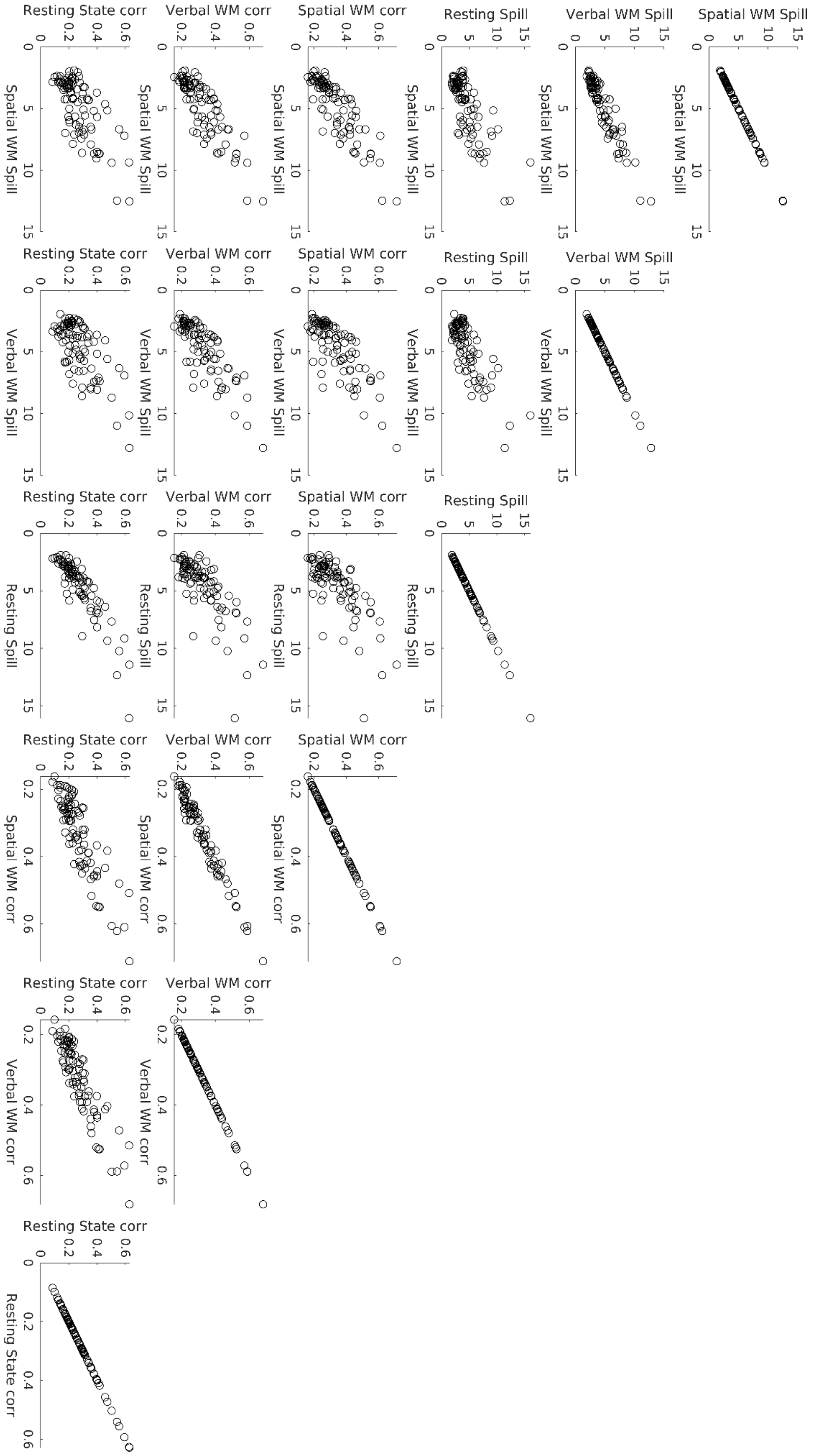


**Conventional connectivity and spillover maps were averaged across individuals. We then correlated the lower triangular sections of the connectivity maps with those of the spillover maps pertaining to the receiving of information. The axes of the plots denote the task and map type (spillover or connectivity).**

**Supplementary Figure 19: Information criteria of VAR models for different lag orders**

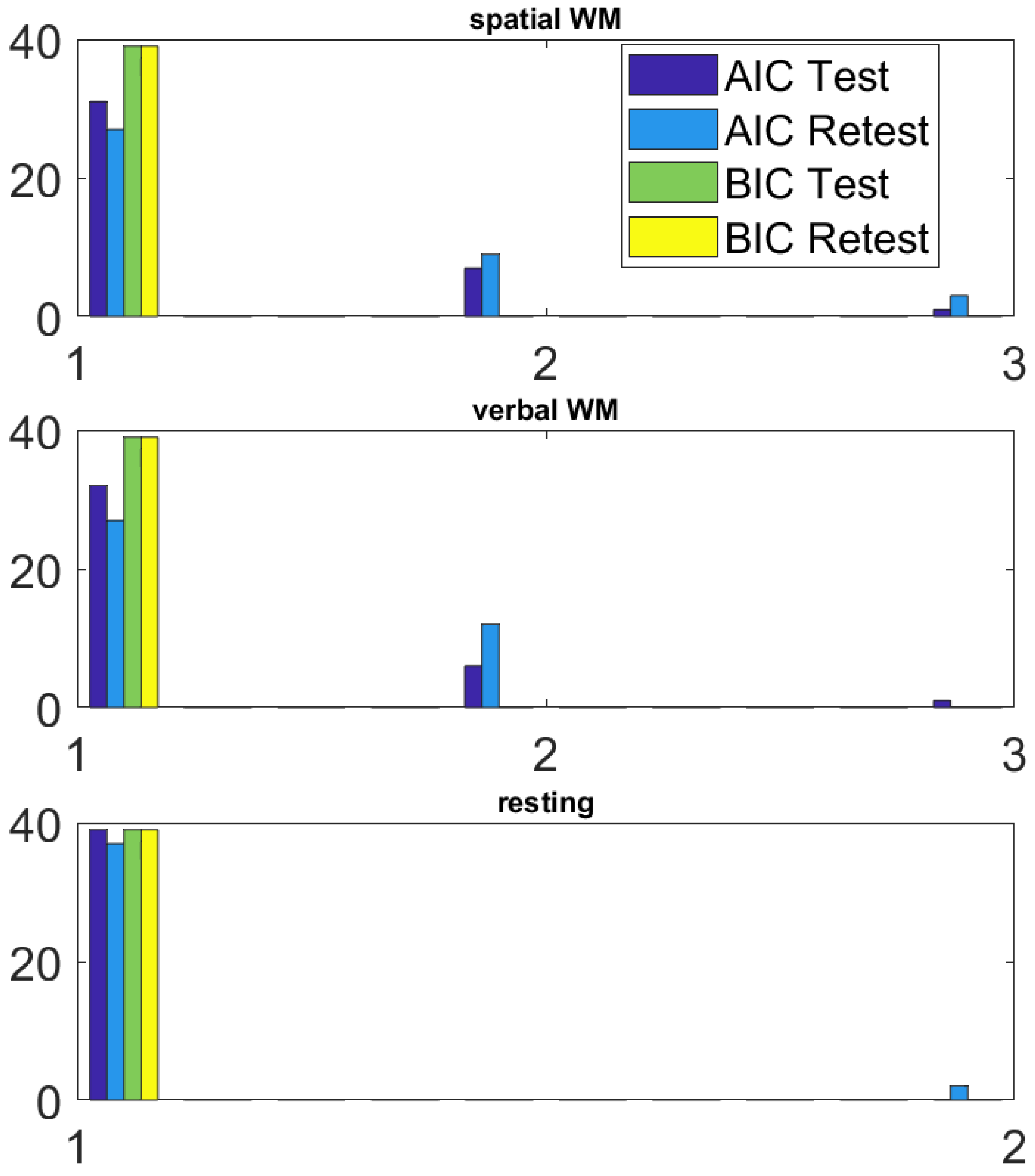


**The 14 timeseries of the 39 individuals were modeled with VAR models of orders 1 to 5. Subsequently the model with the smallest information criterion statistics was selected. The horizontal axis reports the lag that exhibited the best model fit. The overwhelming majority of the individuals showed the best Akaike information criterion (AIC) and Bayesian information criterion (BIC) for 1 lag. In no instances higher lag orders survived the model comparison procedure.**

**Supplementary Figure 20: Residual autocorrelations of VAR models at different lags**

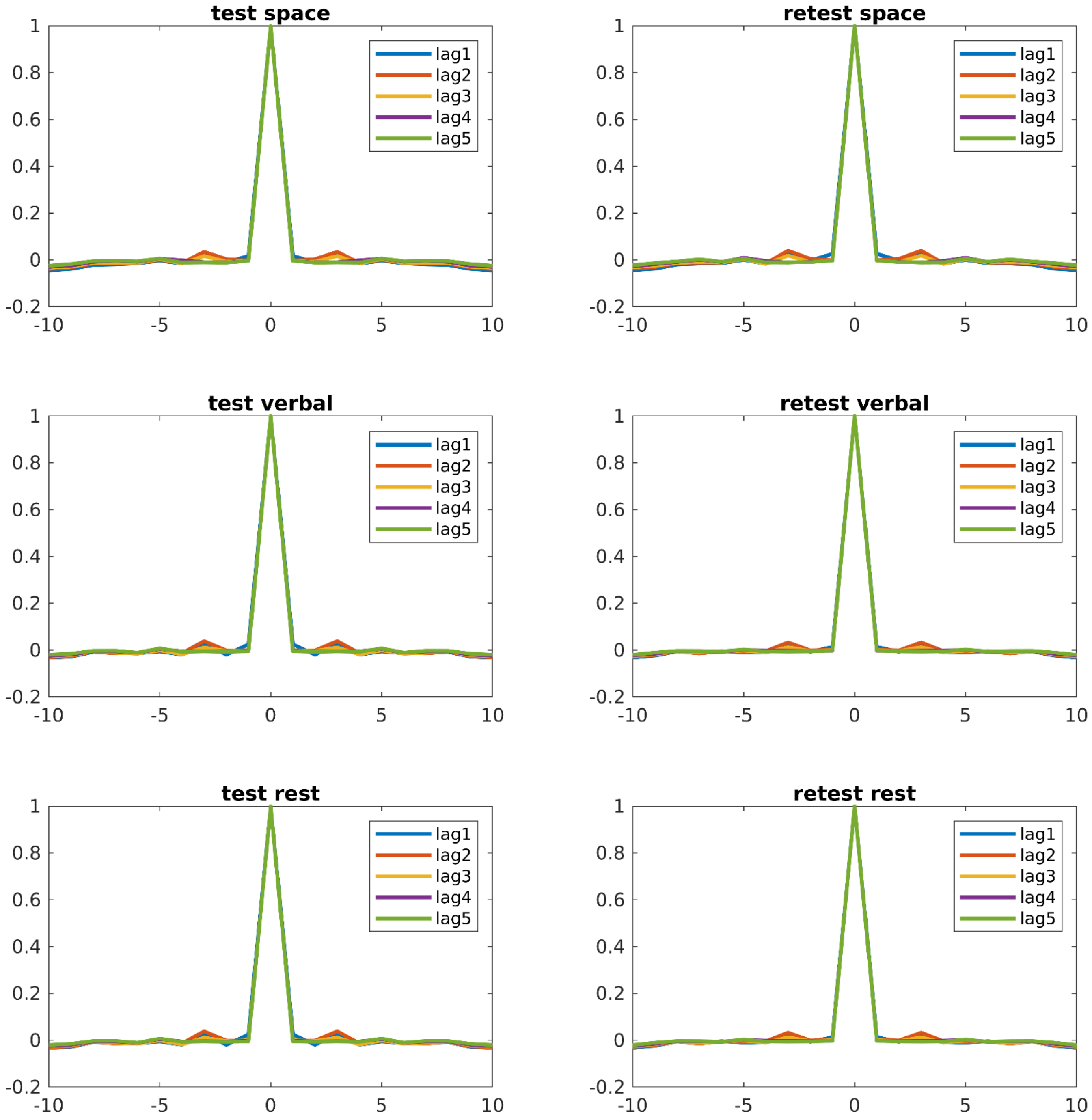


**The residual time course of the VAR models using lag orders 1 to 5 were extracted per individual per ROI per run. Subsequently, we estimated the autocorrelation functions for every VAR model and averaged the results per lag per run. Results show that residuals were uncorrelated attesting the appropriateness of the models.**

### Supplementary Table 1: Correlations of spillover maps across different lag orders

**Space Test**

| | lag 1 | lag 2 | lag 3 | lag 4 |
|---|---|---|---|---|
| lag 2 | 0.9999 | | | |
| lag 3 | 0.9998 | 0.9999 | | |
| lag 4 | 0.9996 | 0.9998 | 0.9999 | |
| lag 5 | 0.9994 | 0.9996 | 0.9997 | 0.9999 |

**Space Retest**

| | lag 1 | lag 2 | lag 3 | lag 4 |
|---|---|---|---|---|
| lag 2 | 0.9998 | | | |
| lag 3 | 0.9996 | 0.9999 | | |
| lag 4 | 0.9994 | 0.9997 | 0.9999 | |
| lag 5 | 0.9993 | 0.9996 | 0.9997 | 0.9999 |

**Space No Diag Test**

| | lag 1 | lag 2 | lag 3 | lag 4 |
|---|---|---|---|---|
| lag 2 | 0.9991 | | | |
| lag 3 | 0.9980 | 0.9993 | | |
| lag 4 | 0.9967 | 0.9983 | 0.9994 | |
| lag 5 | 0.9960 | 0.9973 | 0.9984 | 0.9995 |

**Space No Diag Retest**

| | lag 1 | lag 2 | lag 3 | lag 4 |
|---|---|---|---|---|
| lag 2 | 0.9984 | | | |
| lag 3 | 0.9968 | 0.9993 | | |
| lag 4 | 0.9957 | 0.9985 | 1.000 | |
| lag 5 | 0.9959 | 0.9980 | 0.999 | 1.000 |

**Verbal Test**

| | lag 1 | lag 2 | lag 3 | lag 4 |
|---|---|---|---|---|
| lag 2 | 0.9999 | | | |
| lag 3 | 0.9998 | 1.0000 | | |
| lag 4 | 0.9997 | 0.9999 | 1.0000 | |
| lag 5 | 0.9996 | 0.9998 | 0.9999 | 1.0000 |

**Verbal Retest**

| | lag 1 | lag 2 | lag 3 | lag 4 |
|---|---|---|---|---|
| lag 2 | 0.9999 | | | |
| lag 3 | 0.9998 | 0.9999 | | |
| lag 4 | 0.9997 | 0.9999 | 1.0000 | |
| lag 5 | 0.9997 | 0.9998 | 0.9999 | 1.0000 |

**Verbal No Diag Test**

| | lag 1 | lag 2 | lag 3 | lag 4 |
|---|---|---|---|---|
| lag 2 | 0.9990 | | | |
| lag 3 | 0.9983 | 0.9996 | | |
| lag 4 | 0.9976 | 0.9991 | 0.9997 | |
| lag 5 | 0.9964 | 0.9980 | 0.9989 | 0.9996 |

**Verbal No Diag Retest**

| | lag 1 | lag 2 | lag 3 | lag 4 |
|---|---|---|---|---|
| lag 2 | 0.9989 | | | |
| lag 3 | 0.9985 | 0.9994 | | |
| lag 4 | 0.9979 | 0.9989 | 0.9995 | |
| lag 5 | 0.9972 | 0.9981 | 0.9989 | 0.9996 |

**Rest Test**

| | lag 1 | lag 2 | lag 3 | lag 4 |
|---|---|---|---|---|
| lag 2 | 0.9999 | | | |
| lag 3 | 0.9997 | 0.9999 | | |
| lag 4 | 0.9996 | 0.9998 | 0.9999 | |
| lag 5 | 0.9995 | 0.9997 | 0.9998 | 0.9999 |

**Rest Retest**

| | lag 1 | lag 2 | lag 3 | lag 4 |
|---|---|---|---|---|
| lag 2 | 0.9999 | | | |
| lag 3 | 0.9998 | 0.9999 | | |
| lag 4 | 0.9995 | 0.9996 | 0.9999 | |
| lag 5 | 0.9994 | 0.9995 | 0.9998 | 0.9999 |

**Rest No Diag Test**

| | lag 1 | lag 2 | lag 3 | lag 4 |
|---|---|---|---|---|
| lag 2 | 0.9988 | | | |
| lag 3 | 0.9977 | 0.9990 | | |
| lag 4 | 0.9959 | 0.9976 | 0.9988 | |
| lag 5 | 0.9952 | 0.9970 | 0.9980 | 0.9992 |

**Rest No Diag Retest**

| | lag 1 | lag 2 | lag 3 | lag 4 |
|---|---|---|---|---|
| lag 2 | 0.9988 | | | |
| lag 3 | 0.9975 | 0.9988 | | |
| lag 4 | 0.9961 | 0.9973 | 0.9989 | |
| lag 5 | 0.9953 | 0.9963 | 0.9981 | 0.9990 |

**Space Minus Rest Test**

| | lag 1 | lag 2 | lag 3 | lag 4 |
|---|---|---|---|---|
| lag 2 | 0.9969 | | | |
| lag 3 | 0.9898 | 0.9969 | | |
| lag 4 | 0.9752 | 0.9872 | 0.9951 | |
| lag 5 | 0.9487 | 0.9658 | 0.9795 | 0.9933 |

**Space Minus Rest Retest**

| | lag 1 | lag 2 | lag 3 | lag 4 |
|---|---|---|---|---|
| lag 2 | 0.9961 | | | |
| lag 3 | 0.9893 | 0.9961 | | |
| lag 4 | 0.9719 | 0.9826 | 0.9935 | |
| lag 5 | 0.9474 | 0.9610 | 0.9777 | 0.9939 |

**Space Minus Rest No Diag Test**

| | lag 1 | lag 2 | lag 3 | lag 4 |
|---|---|---|---|---|
| lag 2 | 0.9942 | | | |
| lag 3 | 0.9854 | 0.9951 | | |
| lag 4 | 0.9751 | 0.9880 | 0.9959 | |
| lag 5 | 0.9648 | 0.9795 | 0.9887 | 0.9959 |

**Space Minus Rest No Diag Retest**

| | lag 1 | lag 2 | lag 3 | lag 4 |
|---|---|---|---|---|
| lag 2 | 0.9906 | | | |
| lag 3 | 0.9815 | 0.9949 | | |
| lag 4 | 0.9734 | 0.9886 | 0.9962 | |
| lag 5 | 0.9660 | 0.9823 | 0.9907 | 0.9967 |

**Verbal Minus Rest Test**

| | lag 1 | lag 2 | lag 3 | lag 4 |
|---|---|---|---|---|
| lag 2 | 0.9950 | | | |
| lag 3 | 0.9870 | 0.9961 | | |
| lag 4 | 0.9686 | 0.9838 | 0.9939 | |
| lag 5 | 0.9345 | 0.9561 | 0.9737 | 0.9909 |

**Verbal Minus Rest Retest**

| | lag 1 | lag 2 | lag 3 | lag 4 |
|---|---|---|---|---|
| lag 2 | 0.9966 | | | |
| lag 3 | 0.9891 | 0.9935 | | |
| lag 4 | 0.9672 | 0.9725 | 0.9903 | |
| lag 5 | 0.9302 | 0.9366 | 0.9655 | 0.9896 |

**Verbal Minus Rest No Diag Test**

| | lag 1 | lag 2 | lag 3 | lag 4 |
|---|---|---|---|---|
| lag 2 | 0.9937 | | | |
| lag 3 | 0.9863 | 0.9954 | | |
| lag 4 | 0.9774 | 0.9892 | 0.9959 | |
| lag 5 | 0.9670 | 0.9813 | 0.9895 | 0.9956 |

**Verbal Minus Rest No Diag Retest**

| | lag 1 | lag 2 | lag 3 | lag 4 |
|---|---|---|---|---|
| lag 2 | 0.9938 | | | |
| lag 3 | 0.9879 | 0.9947 | | |
| lag 4 | 0.9812 | 0.9885 | 0.9959 | |
| lag 5 | 0.9730 | 0.9813 | 0.9908 | 0.9953 |

**Space Minus Verbal Test**

| | lag 1 | lag 2 | lag 3 | lag 4 |
|---|---|---|---|---|
| lag 2 | 0.9912 | | | |
| lag 3 | 0.9855 | 0.9947 | | |
| lag 4 | 0.9730 | 0.9847 | 0.9937 | |
| lag 5 | 0.9625 | 0.9753 | 0.9859 | 0.9963 |

**Space Minus Verbal Retest**

| | lag 1 | lag 2 | lag 3 | lag 4 |
|---|---|---|---|---|
| lag 2 | 0.9341 | | | |
| lag 3 | 0.9294 | 0.9380 | | |
| lag 4 | 0.9131 | 0.9226 | 0.9244 | |
| lag 5 | 0.8978 | 0.9058 | 0.9077 | 0.9089 |

**Space Minus Verbal No Diag Test**

| | lag 1 | lag 2 | lag 3 | lag 4 |
|---|---|---|---|---|
| lag 2 | 0.9855 | | | |
| lag 3 | 0.9729 | 0.9893 | | |
| lag 4 | 0.9632 | 0.9766 | 0.9917 | |
| lag 5 | 0.9532 | 0.9661 | 0.9819 | 0.9944 |

**Space Minus Verbal No Diag Retest**

| | lag 1 | lag 2 | lag 3 | lag 4 |
|---|---|---|---|---|
| lag 2 | 0.8637 | | | |
| lag 3 | 0.8580 | 0.8725 | | |
| lag 4 | 0.8325 | 0.8459 | 0.8479 | |
| lag 5 | 0.8128 | 0.8223 | 0.8225 | 0.8294 |

**Spillover maps were estimated for VAR(1) till VAR(5). We averaged the spillover maps across individuals per experimental condition per run per lag. Subsequently, maps were correlated across lags. Due to the fact that the effects of spillover within brain regions were consistently larger than between regions, there is a possibility of artificially inflating the similarity of maps when not accounting for this. Therefore, we chose to calculate across lag correlations both including and excluding within region spillover (No Diag) in order to fully assess the accuracy of our results. The map similarity across lags was very high for all tasks and task contrasts irrespective of analytical approach.**

#### Supplementary  Table 2: Reliability of Net spillover regions

| | *x* | *y* | *z* | *Space* | *Verbal* | *Rest* | *Space-Rest* | *Verbal-Rest* | *Space-Verbal* |
|---|---|---|---|---|---|---|---|---|---|
| *Left AIP* | 34 | -38 | 42 | 0.76 | 0.43 | 0.37 | 0.45 | 0.17 | 0.47 |
| *Left FEF* | 28 | 0 | 58 | 0.76 | 0.72 | 0.37 | 0.59 | 0.59 | 0.29 |
| *Left IFS MFG* | 40 | 16 | 28 | 0.68 | 0.70 | 0.54 | 0.57 | 0.43 | 0.07 |
| *Left LIP* | 40 | -44 | 50 | 0.45 | 0.34 | 0.23 | 0.37 | 0.03 | 0.32 |
| *Left preCG2* | 52 | 2 | 42 | 0.77 | 0.45 | 0.47 | 0.58 | 0.40 | 0.35 |
| *Left Precuneus* | 10 | -54 | 48 | 0.58 | 0.84 | 0.67 | 0.40 | 0.61 | 0.26 |
| *Left STG MTG* | 60 | -50 | 10 | 0.54 | 0.53 | 0.57 | 0.29 | 0.43 | 0.29 |
| *Left VIP1* | 22 | -64 | 54 | 0.56 | 0.44 | 0.35 | 0.44 | 0.34 | 0.50 |
| *Right caudal IPS* | -26 | -66 | 38 | 0.61 | 0.52 | 0.33 | 0.56 | 0.34 | -0.01 |
| *Right IFG tria* | -44 | 18 | 4 | 0.78 | 0.74 | 0.36 | 0.68 | 0.49 | 0.31 |
| *Right LIP* | -36 | -54 | 48 | 0.71 | 0.78 | 0.60 | 0.55 | 0.59 | 0.31 |
| *Right midMFG* | -50 | 32 | 34 | 0.61 | 0.53 | 0.59 | 0.56 | 0.50 | 0.37 |
| *Right precuneus* | -4 | -56 | 54 | 0.59 | 0.51 | 0.49 | 0.45 | 0.41 | 0.39 |
| *Right rostralMFG* | -42 | 46 | 26 | 0.63 | 0.78 | 0.32 | 0.49 | 0.66 | 0.28 |

**This table reports the test-retest reliability estimated with a ICC(2,1) for the net spillover which is estimated by subtracting the "to" or send information from the "from" or received information. The xyz columns display the MNI coordinates while titles above the columns display the experimental condition of the data. Note that complex contrast was made by subtracting one experimental condition from the other before the data were entered into the ICC model.**

#### Supplementary  Table 3: Reliable and significant net spillover regions

| | *x* | *y* | *z* | *Space* | *Verbal* | *Rest* | *Space-Rest* | *Verbal-Rest* | *Space-Verbal* |
|---|---|---|---|---|---|---|---|---|---|
| *Left AIP* | 34 | -38 | 42 | -18.53 | -17.62 | | -19.09 | | |
| *Left FEF* | 28 | 0 | 58 | | | | | | |
| *Left IFS MFG* | 40 | 16 | 28 | 18.40 | | | | | |
| *Left LIP* | 40 | -44 | 50 | -19.54 | | | | | |
| *Left preCG2* | 52 | 2 | 42 | | -13.75 | | | -15.83 | |
| *Left Precuneus* | 10 | -54 | 48 | | | | | -13.93 | |
| *Left STG MTG* | 60 | -50 | 10 | -13.82 | | -11.30 | | | |
| *Left VIP1* | 22 | -64 | 54 | -12.74 | -16.93 | | | | |
| *Right caudal IPS* | -26 | -66 | 38 | | | | | | |
| *Right IFG tria* | -44 | 18 | 4 | | | | 22.78 | 18.19 | |
| *Right LIP* | -36 | -54 | 48 | | | | | | |
| *Right midMFG* | -50 | 32 | 34 | 24.45 | 23.84 | | | 16.56 | |
| *Right precuneus* | -4 | -56 | 54 | 24.02 | 25.21 | 12.99 | | 12.22 | |
| *Right rostralMFG* | -42 | 46 | 26 | | | | | | |

**This table reports the net spillover which survived a Bonferroni corrected conjunction analysis and in addition exhibited sufficient test-retest reliability ICC(2,1)>0.4. The conjunction analysis required that both the test and retest run exhibited significant effects at p =0.05/(14 ROIs * 2). The xyz columns display the MNI coordinates while titles above the columns display the experimental condition and contrasts between experimental conditions.**

#### Supplementary Table 4: Cross spillover map/ connectivity map correlations

| | *Spatial WM Spill* | *Verbal WM Spill* | *Resting Spill* | *Spatial WM Con* | *Verbal WM Con* | *Resting State Con* |
|---|---|---|---|---|---|---|
| *Spatial WM Spill* | 1.00 | 0.98 | 0.81 | 0.93 | 0.92 | 0.81 |
| *Verbal WM Spill* | 0.98 | 1.00 | 0.84 | 0.91 | 0.93 | 0.83 |
| *Resting Spill* | 0.81 | 0.84 | 1.00 | 0.79 | 0.84 | 0.92 |
| *Spatial WM Con* | 0.93 | 0.91 | 0.79 | 1.00 | 0.98 | 0.87 |
| *Verbal WM Con* | 0.92 | 0.93 | 0.84 | 0.98 | 1.00 | 0.90 |
| *Resting State Con* | 0.81 | 0.83 | 0.92 | 0.87 | 0.90 | 1.00 |

| | *Spatial WM Spill* | *Verbal WM Spill* | *Resting Spill* | *Spatial WM Con* | *Verbal WM Con* | *Resting State Con* |
|---|---|---|---|---|---|---|
| *Spatial WM Spill* | 1.00 | 0.95 | 0.76 | 0.87 | 0.86 | 0.72 |
| *Verbal WM Spill* | 0.95 | 1.00 | 0.80 | 0.79 | 0.82 | 0.69 |
| *Resting Spill* | 0.76 | 0.80 | 1.00 | 0.72 | 0.77 | 0.89 |
| *Spatial WM Con* | 0.87 | 0.79 | 0.72 | 1.00 | 0.98 | 0.87 |
| *Verbal WM Con* | 0.86 | 0.82 | 0.77 | 0.98 | 1.00 | 0.90 |
| *Resting State Con* | 0.72 | 0.69 | 0.89 | 0.87 | 0.90 | 1.00 |

**In contrast to symmetrical connectivity maps, spillover maps show directional relations between regions. While connectivity maps have inherent correlations of 1 along the diagonal, spillover maps do not due to varying levels of self-input in regions. Therefore, we omitted the diagonals and correlated the upper and lower triangles of spill over maps with their corresponding connectivity maps. The upper table represents correlations with the "sending" map, while the lower table represents correlations with the “receiving” map. Note that the correlations with the sending map are generally higher when compared to the receiving map.**

#### Supplementary Table 5: Autocorrelation after ARMA(1,1) filter

| | *space* | *verbal* | *rest* |
|---|---|---|---|
| *lag -8* | -0.05 | -0.03 | -0.01 |
| *lag -7* | -0.04 | -0.02 | -0.01 |
| *lag -6* | -0.02 | 0.00 | 0.00 |
| *lag -5* | 0.01 | 0.01 | -0.01 |
| *lag -4* | 0.00 | 0.01 | -0.01 |
| *lag -3* | 0.07 | 0.08 | 0.03 |
| *lag -2* | 0.06 | 0.08 | 0.06 |
| *lag -1* | 0.00 | -0.01 | 0.00 |
| *lag 0* | 1.00 | 1.00 | 1.00 |
| *lag 1* | 0.00 | -0.01 | 0.00 |
| *lag 2* | 0.06 | 0.08 | 0.06 |
| *lag 3* | 0.07 | 0.08 | 0.03 |
| *lag 4* | 0.00 | 0.01 | -0.01 |
| *lag 5* | 0.01 | 0.01 | -0.01 |
| *lag 6* | -0.02 | 0.00 | 0.00 |
| *lag 7* | -0.04 | -0.02 | -0.01 |
| *lag 8* | -0.05 | -0.03 | -0.01 |

**We addressed serial autocorrelations within each time course by applying an ARMA(1,1) filter. Subsequently, we separately estimated the autocorrelation functions for test and retest runs for each subject and each region from lag -8 to lag 8. Finally, we averaged these 39x34x2 autocorrelation functions and presented the results according to their respective task. The low autocorrelations observed suggest that the model that was used to remove autocorrelations was effective.**

#### Supplementary Text 1

A regression analysis conducted on the average resting and working state connectomes showed a strong relation between the two, indicating that the networks involved in working state processes overlap with those involved in resting state while at the same time brain connectivity during working state is somewhat higher as during resting state (**Supplementary Fig. 13**). In analogy to standard connectivity analysis, we conducted a regression analysis on the working state spillover maps and resting state spillover maps. The results displayed in the scatter plot in **Supplementary Fig. 14** demonstrate a strong correlation between the working and resting state maps, similar to standard connectivity analysis (**Supplementary Fig. 13**). It is apparent that the spillover effects within regions are substantially higher than between regions which may bias the observed correlations as within region spillover is outlying in nature. Hence, we created separate scatter plots for the within and between region spillover. The results provided in **Supplementary Fig. 15-16** reveal that the similarities among maps remain robust. We asked ourselves the question whether standard connectivity maps show similarity with spillover maps. A thorough correlation analysis indicates substantial correlations between mean standard connectivity maps and VAR-based maps. It is interesting to note that the correlation between sending spillover maps and connectivity maps is higher than for receiving maps (**Supplementary Fig. Table 4**). The latter was confirmed when scatterplots of the connectivity/spillover relations were inspected (**Supplementary Fig. 17-18**). In conclusion, spillover maps demonstrate a high level of similarity across different lags and tasks, and also exhibit considerable resemblance to standard connectivity maps.

#### Supplementary Text 2

It has been proposed that numbers are represented in a spatial number line, implying that in the number Stroop task, response times may be slower when holding spatial coordinates in memory compared to holding letters in memory. The table below suggest that a small effect of 0.02 seconds might indeed exist.

| | Stroop Spatial | Stroop Verbal | Memory Spatial | Memory Verbal |
|---|---|---|---|---|
| mean RT in seconds | 0.83 | 0.81 | 1.10 | 1.00 |
| percentage wrong responses | 0.37 | 0.59 | 8.28 | 2.56 |
| ICC | 0.78 | 0.88 | 0.72 | 0.74 |

To investigate this, we conducted a conjunction analysis using a one-sided t-test to determine if the number Stroop task with spatial working memory was significantly slower than the task with verbal working memory. The results, with a p-value of 0.77, indicated no such relationship existed in this study. It is commonly believed that the maximum number of items that can be stored in verbal working memory is 7, while for spatial working memory, it is only 4. Thus, the cognitive load for spatial working memory, with two items, should theoretically be higher than for verbal memory with the same number of items. A conjunction analysis showed a p-value of 0.0033, supporting the idea that cognitive load is indeed higher for spatial working memory than for verbal memory. Furthermore, participants made 8.3% mistakes during the spatial WM condition while this was only 2.6% during the verbal WM condition suggesting that the spatial condition is indeed more demanding. The results of test-retest analysis suggest that the absence or presence of the observed difference where truly existent as the ICC(2,1) was in no instance lower than 0.72.
